# Integrated Lineage Tracing in Hepatoblastoma Finds a Plasticity-Proliferation Axis Drives Adaptation

**DOI:** 10.1101/2025.04.16.648783

**Authors:** Emilia Chen, Christie English, Alejandro Allo Anido, Ayeh Sadat Sadr, Marc Miravet-Martí, Laura Zanatto, Andrea Michelle Castaneda Moreno, Siân Hamer, Emilie Inderise, Carolina Armengol, Pau Sancho-Bru, Stefano Cairo, Alejandra Bruna

**Affiliations:** Centre for Evolution and Cancer, Division of Cancer Biology, The Institute of Cancer Research (ICR), London, UK; Institut d’Investigacions Biomèdiques August Pi i Sunyer (IDIBAPS), Barcelona, Spain; Institute for Health Science Research Germans Trias i Pujol (IGTP), Badalona, Spain; University of Barcelona (UB), Barcelona, Spain; Biomedical Research Networking Centre in Hepatic and Digestive Diseases (CIBEREHD), Madrid, Spain; XenTech, Évry-Courcouronnes, France; Fondazione Istituto di Ricerca Pediatrica Città della Speranza (IRP), Padova, Italy; Champions Oncology, Inc., Hackensack, New Jersey, USA

## Abstract

Paediatric cancers, with their developmental context and low mutational burden, are powerful models for studying plasticity in cancer evolution. Hepatoblastoma (HB) has among the lowest mutational burdens of childhood cancers, yet ∼20% relapse, implicating non-genetic mechanisms in treatment adaptation. We integrate expressed DNA barcodes with longitudinal single-cell multiomics to trace clonal and phenotypic dynamics following cisplatin treatment in HB cell lines. Cisplatin imposes a phenotypic bottleneck, selecting for progenitor-like cells across clones without a clonal sweep. Exit from persistence involves asynchronous, partly fitness-biased re-expansion and re-diversification towards hepatocytic-like states, also captured in two longitudinal patient cases. Pseudotime and dynamical-landscape analyses reveal a plasticity-proliferation axis underlying recovery, where persister cells occupy an arrested-plastic state, exited to restart proliferation. Chromatin and regulatory analyses identify the E2F target *BIRC5* (survivin) as a key node in this axis, epigenetically primed post-recovery. *BIRC5* maintains hepatocytic-like states, and its knockdown shifts cells toward progenitor-like phenotypes, while pharmacological inhibition selectively eliminates cisplatin-tolerant persisters. These findings position *BIRC5* as a regulator of plasticity-driven adaptation and a candidate vulnerability in the minimal-residual disease window.

## Introduction

Paediatric cancers have lower mutational burdens than adult cancers, with hepatoblastoma (HB) having one of the lowest frequencies of somatic coding mutations among paediatric cancer types (Gröbner *et al*, 2018). The vast majority of HBs carry mutations that affect the Wnt pathway, with 80% of HBs carrying mutations in *CTNNB1* that activate β-catenin (Koch *et al*, 1999; Eichenmüller *et al*, 2014). The paucity of driver mutations beyond β-catenin activating alterations (Hirsch *et al*, 2021; Nagae *et al*, 2021), combined with the developmental context in which these tumours arise, makes HB an interesting model for studying the role of phenotypic plasticity in cancer evolution.

HB is the most common liver cancer in young children, typically developing within the first five years of life on nonfibrotic liver (Darbari, 2003). Although rare, occurring in approximately one per million children per year, in recent years the incidence rate has been increasing (Hubbard *et al*, 2019). HB treatment follows a cisplatin-based chemotherapy regime followed by surgery. This achieves a 5-year survival rate of close to 80%, however treatment options for chemoresistant tumours remain limited (Feng *et al*, 2019; Wang *et al*, 2024), resulting in poor prognosis(Trobaugh-Lotrario *et al*, 2017).

Despite a lack of genetic driver mutations, HB tumours are phenotypically heterogeneous. HBs have been previously split into two subtypes: C1 and C2 (Cairo *et al*, 2008). Although both have a similar *CTNNB1* mutation profile, their gene expression landscapes are distinct; the C1 subtype corresponds to well-differentiated foetal histotypes and a better prognosis, while C2 subtypes typically exhibit immature histotypes and a worse prognosis. Further stratification of these transcriptomic subtypes (Hirsch *et al*, 2021; Hooks *et al*, 2018; Song *et al*, 2022; Sumazin *et al*, 2022), as well as incorporation of epigenetic profiling (Carrillo-Reixach *et al*, 2020b; Sekiguchi *et al*, 2020), has since expanded on this classification. The heterogeneity in this developmental tumour, driven by few mutations, alludes to non-genetic sources of variation playing a central role in cancer evolutionary processes, such as adaptation to chemotherapy.

Plasticity enables the expression of different phenotypes by the same genotype, under different environmental conditions (Schlichting & Pigliucci, 1998; West-Eberhard, 2003). Across evolutionary and developmental biology, adaptive plasticity has now has a well-established role in creating diversity and facilitating genetic evolution (Sommer, 2020; Price *et al*, 2003). This notion has permeated the study of cancer, garnering appreciation that the hijacking of phenotypic plasticity by cancer cells mediates tumour progression and the development of chemoresistance (Gupta *et al*, 2019; Yuan *et al*, 2019). Although more widely studied in adult cancers, there is growing evidence that plasticity has a role in HB (Song *et al*, 2022; Lee-Theilen *et al*, 2022; Huang *et al*, 2022; Roehrig *et al*, 2024b). However, explicit attention at temporal and molecular resolution to the post-treatment period, during which drug-tolerant persistent cells regrow (persistence-to-recovery), is lacking.

In this study, we combine single-cell RNA sequencing (scRNA-seq) and single-nuclei multiome sequencing (snRNA-seq and snATAC-seq) with expressible DNA static barcodes in HB models to track phenotypic transitions and clonal dynamics in the early stages of adaptive evolution. Through this, we add to the characterisation of plasticity in paediatric cancers and explore plasticity drivers that play a role in the early persistence-to-recovery stage of treatment adaptation, thus serving as prospective therapeutic targets.

## Results

### Cisplatin selects for progenitor-like persisters that fuel re-diversification during recovery

To dissect how phenotypic plasticity drives adaptation to chemotherapy in HB at clonal resolution, we combined expressible DNA barcoding with scRNA-seq and cisplatin treatment in two complementary HB preclinical models: HuH6 and HepG2. Stably barcoded cell line populations are particularly suited to this question, as they enable tracking of large numbers of single-cell-derived clones across longitudinally sampled treatment phases, an approach not yet feasible in primary tumour material at the temporal resolution required to interrogate persistence-to-recovery dynamics.

Since their respective derivations in the 1970s (Doi, 1976; López-Terrada *et al*, 2009; Aden *et al*, 1979), the HuH6 and HepG2 cell lines have become widely used models to study childhood liver cancer. HuH6 and HepG2 were derived from patients of different ages (1 and 15 years, respectively) and differ in oncogenic context. Both exhibit constitutive activity of the Wnt pathway – HuH6 through a *CTNNB1* G34V missense mutation in exon 3 and HepG2 via a large deletion spanning exon 3 and exon 4 (Koch *et al*, 1999) – and functional p53 activity (Hsu *et al*, 1993; Yamamoto *et al*, 2007). Additionally, HepG2 notably harbours a *TERT* mutation (Cevik *et al*, 2015), characteristic of ‘tween’ HB (Nagae *et al*, 2021). Thus, together these cell lines provide a basis to compare conserved versus context-specific cisplatin response dynamics.

Barcoded cells were treated with cisplatin (IC50) for 48 hours, after which a small fraction of cells tolerated and persisted through treatment (persistence phase); following cisplatin removal, these persister cells underwent transcriptomic changes (awakening phase) and eventually resumed proliferation at baseline rates, regenerating cultures morphologically indistinguishable from pre-treatment populations (recovery phase). Longitudinal sampling for scRNA- and DNA sequencing across these phases enabled us to characterise phenotypic and clonal dynamics (Figure 1A; Figure EV1A, B; Appendix Table S1).

**Figure 1:**
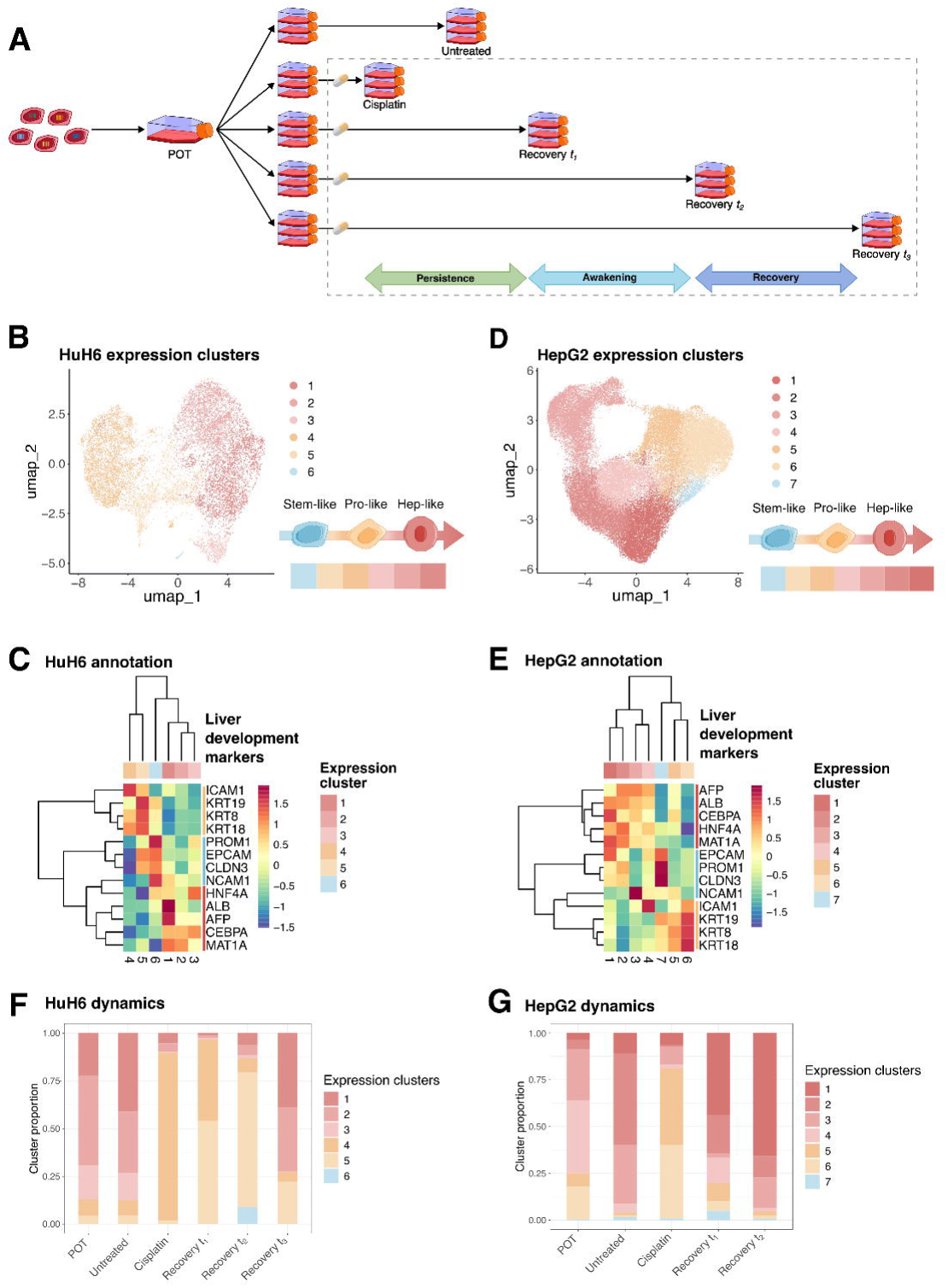
**(A)** Integrated DNA barcoding and single-cell sequencing experiment design. Cells were dosed with cisplatin (IC50) for 48 hours and left to recover until they were observed to be doubling at their baseline proliferation rates (see Methods). **(B)** Uniform manifold approximation and projection (UMAP) of 10,791 HuH6 cells that passed quality control (QC), coloured by numbered expression cluster. Schematic representation of relation between HuH6 expression clusters and liver developmental states. **(C)** Expression of canonical liver development markers (Lemaigre, 2009; Si-Tayeb *et al*, 2010; Schmelzer *et al*, 2007; So *et al*, 2020; Ramani & Lu, 2017) in HuH6 scRNA-seq data, pseudobulked by cluster. *PROM1*, *CLDN3*, *NCAM1* are hepatic stem cell markers; *EPCAM*, *KRT8*, *KRT18* and *KRT19* are hepatic stem cell and hepatoblast markers; *ICAM1* is a hepatoblast marker; *HNF4A* has a role in hepatocyte differentiation; *AFP* is a foetal hepatocyte marker; *CEBPA* and *ALB* are hepatocyte cell fate markers; *MAT1A* is expressed in the adult liver. **(D)** UMAP of 43,017 HepG2 cells that passed QC, coloured by annotated expression cluster. Schematic representation of relation between HepG2 expression clusters and liver developmental states. **(E)** Expression of canonical liver development markers (Lemaigre, 2009; Si-Tayeb *et al*, 2010; Schmelzer *et al*, 2007; So *et al*, 2020; Ramani & Lu, 2017) in HepG2 scRNA-seq data, pseudobulked by cluster. **(F)** Bar chart of HuH6 expression cluster proportions at each experimental timepoint. Colour indicates expression cluster. **(G)** Bar chart of HepG2 expression cluster proportions at each experimental timepoint. Colour indicates expression cluster.

To annotate cell states in our scRNA-seq data, we projected published HB transcriptomic subtypes – Cairo C1/C2 (Cairo *et al*, 2008), Carrillo-Reixach Epi-CA/CB and 14q32 (Carrillo-Reixach *et al*, 2020b), Roehrig sc-H/LP/M/epi (Roehrig *et al*, 2024b). At the model level, both HuH6 and HepG2 fell within a Cairo C2-like, Roehrig liver progenitor (LP)-like space, consistent with their cell line provenance (Figure EV1C-E). Within each model however, these signatures were expressed relatively uniformly across clusters and did not sufficiently resolve intra-model heterogeneity (Figure EV1C, D – hepatoblastoma signatures). Because our question concerns plasticity dynamics that operate at a finer scale than HB subtype classification, we also applied a developmentally informed canonical liver development marker annotation, predicting that phenotypic changes within a given tumour might follow the developmental trajectories of the originating embryonal tissue (Lemaigre, 2009; Si-Tayeb *et al*, 2010; Schmelzer *et al*, 2007; So *et al*, 2020; Ramani & Lu, 2017) (Figure 1C, E; Appendix Table S2). The C2-fixed background shared by both models is a feature of this design: against a stable background subtype, our annotation resolved model-specific, treatment-related phenotypic shifts. Subsequent changes in Cairo C2 signature during cisplatin recovery could be relative shifts within a fixed subtype or proliferation dynamics rather than subtype changes.

Clustering of 10,791 HuH6 single-cell transcriptomes revealed six expression clusters (1-6), which we mapped onto three developmentally defined states (Hep-/Pro-/Stem-like) (Figure 1B). We annotated the clusters predicted in our data using HB signatures (Cairo *et al*, 2008; Carrillo-Reixach *et al*, 2020b; Roehrig *et al*, 2024b), as well as liver development signatures (Wesley *et al*, 2022) and markers (Lemaigre, 2009; Si-Tayeb *et al*, 2010; Schmelzer *et al*, 2007; So *et al*, 2020; Ramani & Lu, 2017) (Figure 1C; Figure EV1C). Hep-like states (clusters 1-3) were enriched for Cairo C2 signature, strongly expressed the Wesley hepatocyte signature and hepatocyte markers (*ALB*, *AFP*, *CEBPA*, *MAT1A*, *HNF4A*). Pro-like states (clusters 4-5) co-expressed Wesley hepatocyte and cholangiocyte signatures and expressed canonical hepatoblast markers (*ICAM1*, *KRT8*, *KRT18*, *KRT19*), consistent with a more bipotent, hepatoblast-like identity. Cluster 6 additionally expressed hepatic stem cell markers (*PROM1*, *EPCAM*, *CLDN3*, *NCAM1*) and defined a Stem-like state, a transient population that, as detailed below, was enriched only during the awakening phase post treatment and given its small size may not represent a stable level in a hierarchy of states.

Analogous analysis of HepG2 cells (Figure 1D, E; Figure EV1D) identified seven expression clusters (1-7) that resolved into the same Hep-/Pro-/Stem-like states defined in the HuH6 data (Figure EV1F), supporting a degree of conservation of this developmental annotation across these HB cell lines. However, the marker patterns distinguishing each state at the cluster level showed heterogeneity between models, consistent with the distinct developmental origins of HuH6 and HepG2 producing model-specific cluster-level structure on a shared differentiation-related cell state axis.

Tracking phenotypic composition along the persistence-awakening-recovery drug response course revealed conserved selection then re-diversification dynamics in both models (Figure 1F, G). In pre-treatment populations, the Hep-like state dominated and the Pro-like state was present at a lower frequency. Cisplatin treatment selected for the Pro-like state, consistent with a phenotypic bottleneck at persistence. During the awakening phase there was also a transient, relative enrichment of the Stem-like population before recovery regenerated the Hep-like dominance characteristic of pre-treatment populations. Across both models, this pattern positions Pro-like states as reservoirs that sustain recovery via plasticity-led regeneration of Hep-like dominance.

### Barcode tracing reveals a lineage-independent phenotype bottleneck resolved by plasticity-led clonal re-expansion

To study clonal dynamics in parallel with phenotypic dynamics during the cisplatin response, we paired our scRNA-seq experiment with expressible DNA barcoding in HuH6 and HepG2. The CloneTracker™ lentiviral barcoding approach introduces a synthetic, heritable DNA sequence per cell that acts as a lineage tag, allowing individual cell lineages to be tracked across treatment phases. Because the sequence is expressed, it is detected in scRNA-seq independently of genetic diversity, making the approach particularly suited to interrogating plasticity dynamics – especially valuable here, where we use genetically stable and uniform preclinical cancer cell populations. One clone is identified by one expressed DNA barcode and represents the founder cell and all its lineage descendants, which share the same barcode. To enable cross-replicate comparison, the barcoded founder population was first expanded so that each founder barcode was represented by multiple daughter cells (the ‘POT’ pool). Seeding from the POT into multiple flasks produced replicates that shared the same set and frequencies of barcoded lineages, allowing each clone to be tracked across treatment conditions and replicates (Acar *et al*, 2020; Roux *et al*, 2023).

During the persistence-awakening-recovery phases, we reasoned that two key evolutionary pressures select for cells that (i) survive cisplatin treatment and (ii) can resume proliferation. While scRNA-seq of barcoded HuH6 cells showed that cisplatin treatment enriched for Pro-like states (indicating their greater fitness under treatment), at the clonal level we did not observe strong clonal sweeps across replicates (at the Cisplatin timepoint; Figure 2A). This is further illustrated by beta diversity (Bray-Curtis dissimilarity; Figure 2B): clonal composition was highly similar across replicates at the Untreated, Cisplatin, Recovery t_1_ and Recovery t_2_ timepoints – indicating no detectable clonal fitness differences, in contrast to the phenotypic space. By contrast, at Recovery t_3_, the replicates diverged markedly from one another and from earlier phases. Proliferation restart occurred with similar timing across replicates, indicating a common recovery window; within that window, the identity of re-proliferating clones differed between replicates. Although the proportion of cells per clone varied, a reproducible fitness signal was present in a subset of clones (Figure 2C): the most abundant clones at Recovery t_3_ in one replicate tended also to be more abundant in others (e.g. Barcode1662, Barcode5495, Barcode6423), while low-abundance clones remained low across replicates (e.g. Barcode1638, Barcode9953, Barcode34070). These observations are most consistent with asynchronous clonal awakening, in which persister clones resume proliferation at different rates within the recovery window, partly stochastically and partly biased by inter-clonal fitness differences. From this data we cannot determine whether all persister clones retain latent proliferative potential, or whether only a subset are competent to resume. Overall, cisplatin selects for a Pro-like persister state but does not impose a strong clonal bottleneck; exit from persistence proceeds through asynchronous, partly fitness-biased clonal re-expansion.

**Figure 2:**
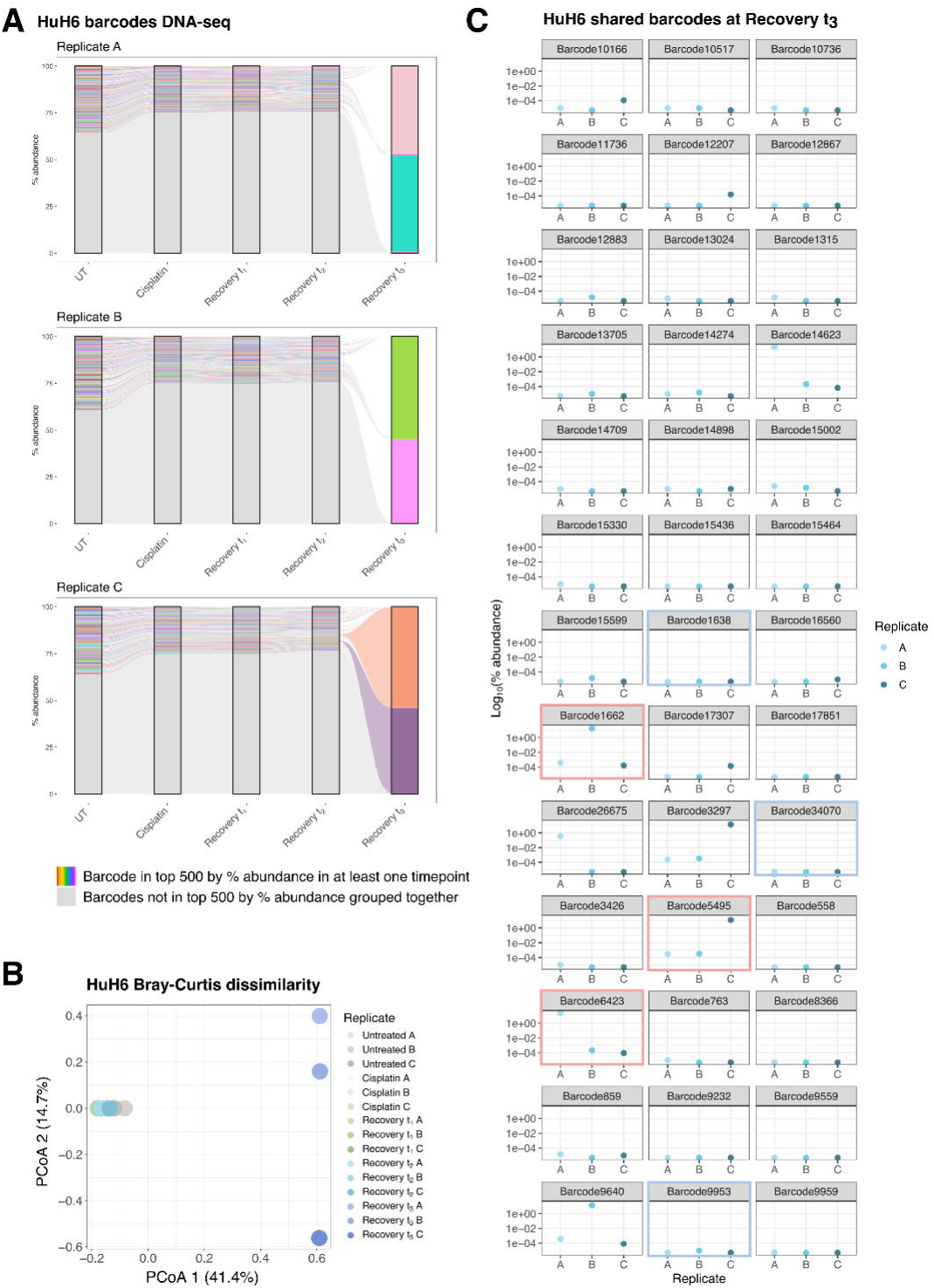
**(A)** Percentage abundance of a sample of unique DNA barcodes tracked over persistence, awakening and recovery phases. Each graph represents one replicate (A, B, C). In each replicate graph, each coloured stratum is an individual barcode, and flows connect the same clone between consecutive timepoints. Each clone has the same colour in every timepoint and replicate. For each timepoint-sample, the 500 most abundant barcodes are shown individually, and all remaining barcodes are combined into a single grey stratum. Because the top 500 barcodes were selected separately for each timepoint-sample, a flow that begins or ends between timepoints shows that the clone entered or left that sample’s top 500; it does not mean the clone was absent. **(B)** Principal coordinate analysis (PCoA) plot of the beta diversity, based on the Bray-Curtis dissimilarity, of the DNA barcode populations between replicates. **(C)** Log_10_ percentage abundance of HuH6 DNA barcodes shared between all three replicates at Recovery t_3_. Each panel is one barcode. Red highlighted panels were used as examples in the main text as examples of expanded barcodes; blue highlighted panels were used as examples in the main text as lower frequency barcodes.

We next integrated expressed barcode information with scRNA-seq profiles to directly link clonal identity to cell state (Appendix Figure S1). In the HuH6 integrated data, most clones already displayed phenotypic heterogeneity prior to cisplatin treatment, indicating innate plasticity; despite each clone originating from a single barcoded cell, by the Untreated timepoint most clones spanned multiple states, showing that plasticity generates pre-existing intra-clonal heterogeneity (Figure 3A). These observations were reproduced in the HepG2 integrated data (Figure 3B).

**Figure 3:**
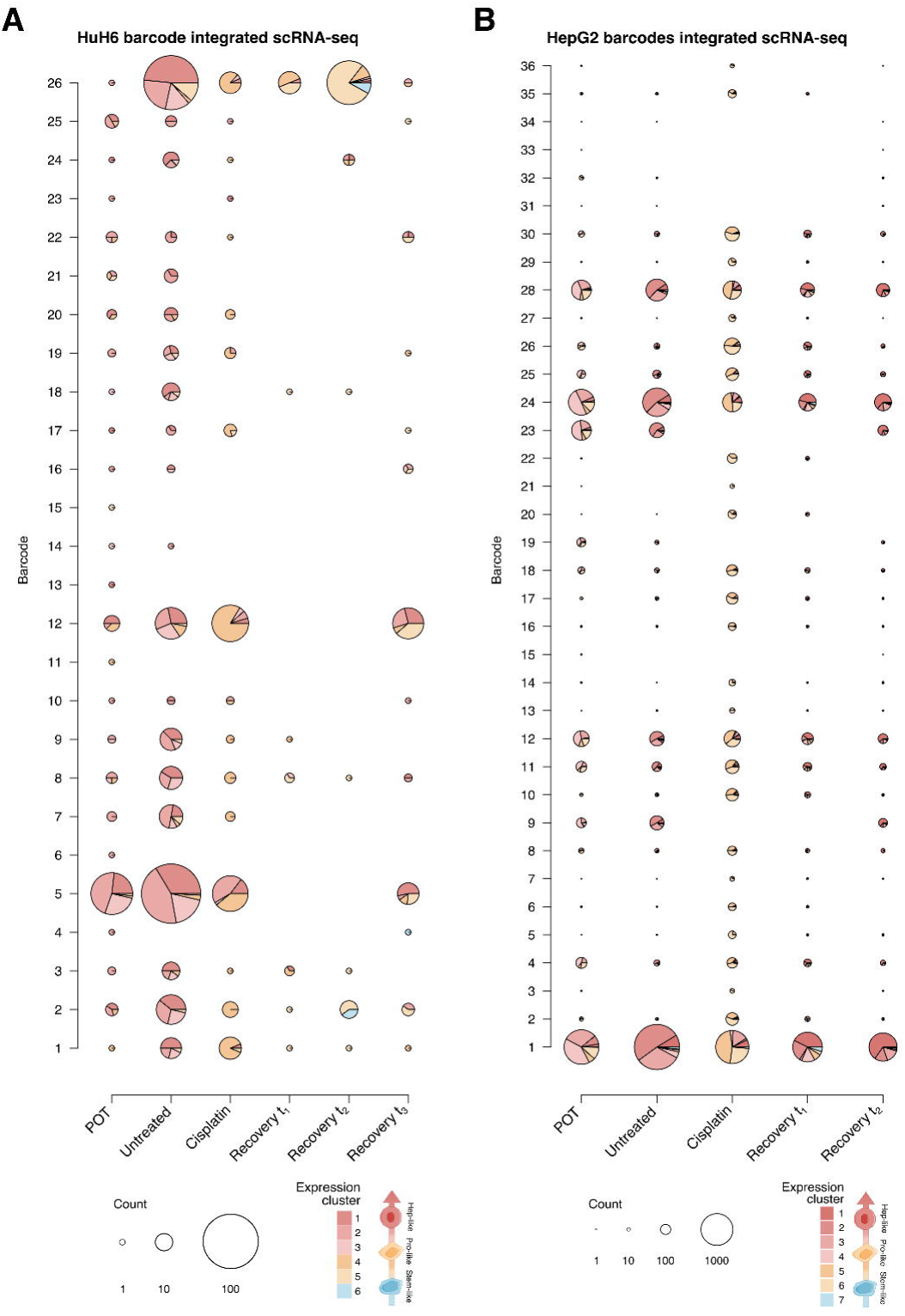
**(A)** Compound bubble-pie chart for HuH6 barcode-integrated scRNA-seq data of 26 expressed DNA barcodes (rows) at each experimental timepoint (columns). The size of the bubble-pie represents the numbers of cells with a given barcode at a given experimental timepoint. The relative sector size for each bubble-pie represents the proportion of cells belonging to an expression cluster (colour), for a given barcode at a given experimental timepoint. *N.B.*: Barcode numbers were reassigned in the integrated data so do not relate to those in Figure 2. **(B)** Compound bubble-pie chart for HepG2 barcode-integrated scRNA-seq data of 36 expressed DNA barcodes (rows) at each experimental timepoint (columns). The size of the bubble-pie represents the numbers of cells with a given barcode at a given experimental timepoint. The relative sector size for each bubble-pie represents the proportion of cells belonging to an expression cluster (colour), for a given barcode at a given experimental timepoint.

Following treatment, clones were enriched for the Pro-like state, and during recovery they regained the phenotypic diversity of the pre-treatment population. Lineage-resolved longitudinal analysis showed that the phenotypic dynamics of most clones paralleled the population-level pattern: an initially heterogeneous pool dominated by the Hep-like state shifted towards Pro-like at persistence and regained pre-treatment heterogeneity through recovery. A subset of clones additionally underwent transient phenotypic sweeps, consistent with intra-clonal plasticity. For example, in the HuH6 experiment the population of clone 2 consisted exclusively of Pro-like cluster 4 during the persistence phase, transitioned to Pro-like cluster 5 and Stem-like cluster 6 during awakening, before the Hep-like state re-emerged after recovery. It should be noted however, that in general, phenotypic shifts were moderate and not complete sweeps. Together, these dynamics indicate a lineage-independent phenotypic bottleneck in response to cisplatin and that plasticity from Pro-like to Hep-like states drives exit from persistence and underpins post-treatment regrowth both at the population level and at single clone resolution. Recovery thus integrates phenotypic selection at persistence with stochastic, plasticity-led re-expansion at awakening, regenerating population heterogeneity after treatment.

### Longitudinal clinical case studies in two HB subtypes show shared plasticity dynamics during treatment response

To examine whether the persistence-to-recovery dynamics observed in our preclinical models were also evident in HB tumours, we performed longitudinal single-nuclei RNA sequencing (snRNA-seq) on tumour samples from two patients (P1 and P2) with HB. Each patient was sampled at an earlier (T1, during or shortly after treatment), and a later post-treatment timepoint (T2) (Table 1; Appendix Table S3). The two cases differed in HB subtype context: P1 samples were of a C1-like tumour with foetal epithelial histology, and P2 samples were of a C2-like tumour with mixed epithelial and mesenchymal histology, in which the T2 sample was a lymph node metastasis. This subtype contrast allowed us to ask whether shared plasticity dynamics operate across HB subtypes despite their distinct molecular and histological backgrounds. In the snRNA-seq data, tumour clusters were annotated using HB signatures from literature (Cairo *et al*, 2008; Carrillo-Reixach *et al*, 2020b; Roehrig *et al*, 2024b), liver development signatures (Wesley *et al*, 2022) and canonical liver development markers (Lemaigre, 2009; Si-Tayeb *et al*, 2010; Schmelzer *et al*, 2007; So *et al*, 2020; Ramani & Lu, 2017), which showed that clusters expressed a mixture of markers, consistent with the known phenotypic heterogeneity observed in patient cancer samples (Figure 4A-D; Figure EV2A, B). We also annotated cell types using SingleR with the Human Cell Atlas as the reference and used InferCNV to confirm the identity of tumour cells (Figure EV2C-F; Appendix Table S4).

**Figure 4:**
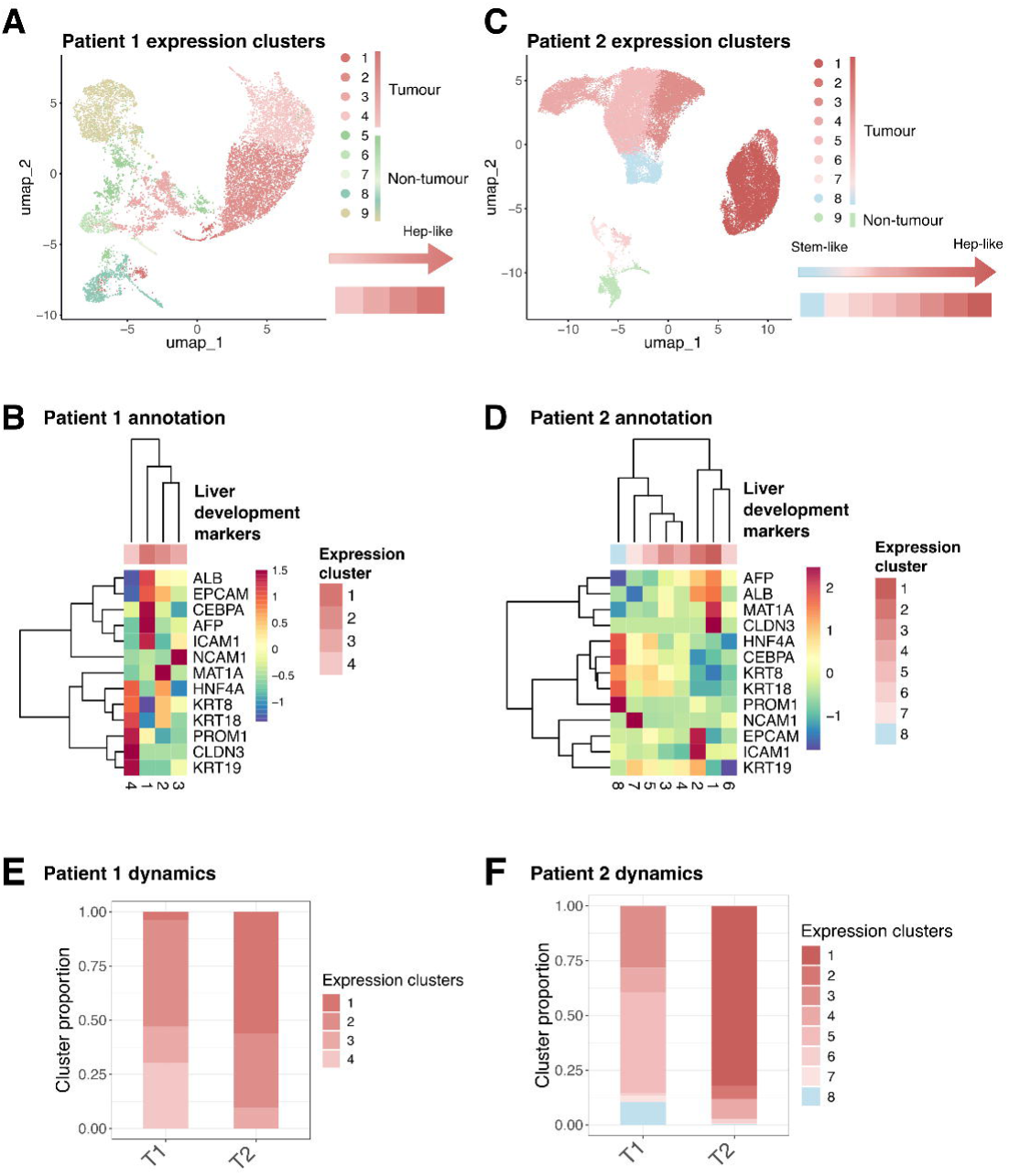
**(A)** Patient 1 (P1) UMAP of 11,858 cells, coloured by expression clusters. Schematic representation of relation between P1 expression clusters and liver developmental states. **(B)** Expression of canonical liver development markers (Lemaigre, 2009; Si-Tayeb *et al*, 2010; Schmelzer *et al*, 2007; So *et al*, 2020; Ramani & Lu, 2017) in P1 snRNA-seq data, pseudobulked by cluster for tumour clusters only (1-4). **(C)** Patient 2 (P2) UMAP of 37,158 cells, coloured by expression clusters. Schematic representation of relation between P2 expression clusters and liver developmental states. **(D)** Expression of canonical liver development markers (Lemaigre, 2009; Si-Tayeb *et al*, 2010; Schmelzer *et al*, 2007; So *et al*, 2020; Ramani & Lu, 2017) in P2 snRNA-seq data, pseudobulked by cluster for tumour clusters only (1-8). **(E)** Proportion of each P1 cluster in the tumour cell population at T1 (mid-treatment) and T2 (post-treatment relapse). **(F)** Proportion of each P2 cluster in the tumour cell population at T1 (post-treatment) and T2 (post-treatment relapse).

P1 samples were from a C1-like tumour with epithelial foetal histology; both T1 (mid-treatment) and T2 (post-treatment) samples were taken from the liver. snRNA-seq analysis identified nine expression clusters (1-9) of which clusters 1-4 were tumour (supported by InferCNV) and dominated by hepatocyte cell type annotation, while clusters 5-9 were stromal and immune (Figure 4A; Figure EV2C, E). All tumour clusters (1-4) expressed the Cairo C1 signature and were enriched for Wesley hepatocyte signature (Figure EV2A). Within this C1 background, clusters resolved along a hepatocyte-to-hepatoblast differentiation gradient: cluster 1 strongly expressed hepatocyte markers (*ALB*, *MAT1A* and *HNF4A*) with mild progenitor marker expression (*ICAM1, EPCAM*); cluster 2 expressed hepatocyte markers (*ALB*, *CEBPA*, *MAT1A, HNF4A*) alongside progenitor markers (*KRT8*, *KRT18*); cluster 3 expressed lower *ALB* and *AFP* with stronger *ICAM1* and *NCAM1*; and cluster 4 expressed hepatoblast markers (*KRT8*, *KRT18*, *KRT19*) and hepatic stem cell markers (*CLDN3* and *PROM1*) together with the hepatocyte fate marker *HNF4A*, but lacked *ALB* or *AFP* expression (Figure 4B). The four tumour clusters thus span a hepatocytic-to-hepatoblast-like gradient within a single C1-subtype tumour.

P2 samples were of a C2-like tumour with mixed epithelial-mesenchymal histology. Both samples were taken post treatment: T1 from the primary liver tumour (predominantly epithelial) and T2 from a lymph node metastasis. Because T2 represents a different anatomical site, the P2 comparison between T1 and T2 carries an additional caveat that biopsy site differences may contribute to observed changes alongside genuine post-treatment dynamics. snRNA-seq analysis identified nine expression clusters (1-9): clusters 1-8 were tumour (supported by InferCNV) and dominated by hepatocytes in cell type annotation, while cluster 9 was stroma (Figure 4C; Figure EV2D, F). The majority of tumour clusters expressed the Cairo C2 signature and were enriched for the Wesley hepatocyte signature (Figure EV2B). Cluster 8 was an exception: it expressed the Cairo C1 signature and co-expressed the Wesley hepatocyte and cholangiocyte signatures, characteristic of a bipotent progenitor-like population. Within this C2 background, tumour clusters resolved along a hepatocytic-to-progenitor gradient: clusters 1 and 2 were the most hepatocyte-like, expression the highest *ALB* and *AFP* alongside expression of some progenitor markers (*EPCAM*, *CLDN3*, *ICAM1*); clusters 3-7 expressed *ALB*, *AFP* and *MAT1A* at lower levels, with a subset (3-5, 7) expressing hepatocyte fate markers (*HNF4A*, *CEBPA*) together with hepatoblast markers (*KRT8*, *KRT18*); cluster 8 also expressed *HNF4A*, *CEBPA*, *KRT8* and *KRT18*, as well as the progenitor marker *PROM1*, and expressed relatively low levels of *ALB*, *AFP* and *MAT1A* (Figure 4D). The eight tumour clusters thus span hepatocytic-like through hepatoblast-like to a less-differentiated, bipotent progenitor-like state (cluster 8) within a single C2-subtype tumour.

Despite their distinct subtype contexts, P1 and P2 exhibited the same direction of temporal, developmental-like cell state dynamics as those observed in our cell lines. In P1, cluster 4 (the most hepatoblast-like, with *KRT*/*CLDN3*/*PROM1* expression) was present at T1 but lost at T2, while cluster 1 (the most-hepatocytic-like, with strongest *ALB* and *AFP* expression) became dominant at T2 (Figure 4E). In P2, the less-differentiated, bipotent progenitor-like cluster 8 was present at T1 but its proportion shrunk substantially at T2, while cluster 1 (the most-hepatocyte-like, with the strongest *ALB* and *AFP* expression) dominated at T2 (Figure 4F). Although this is a case study of two patients, since we were limited by the availability of longitudinal samples, both patients show shrinkage of less-differentiated tumour clusters and expansion of the most hepatocyte-like cluster between T1 and T2, paralleling the Pro-to-Hep direction observed in cell lines across the awakening-to-recovery post treatment window. Notably, the cell states traversed during these trajectories differ slightly by subtype: P1 (C1-like) moves along a hepatocyte-to-hepatoblast axis, whereas P2 (C2-like) additionally engages a less-differentiated, progenitor-like state (cluster 8). These trends also mirror observations in scRNA-seq data of 5 patients, pre- and post-chemotherapy from Chen *et al*. (2025, 2026) (Appendix Figure S2): three out of the five patients show post-treatment enrichment of hepatocyte-associated genes (*CEBPA*, *HNF4A*, *ALB*, *MAT1A*) and exhibited a “partial response”, while the two patients that were not enriched post-treatment for hepatocyte-associated gene expression displayed “stable disease”. Together, this supports the two cases we described, indicating that an analogous Pro-to-Hep plasticity operates across HB subtypes post treatment, though the phenotypic space it occupies is shaped by subtype context.

### Pseudotemporal trajectories identify coupling of plasticity and proliferation programs during recovery

To complement our longitudinal barcoding approach, we used an ensemble pseudotime analysis, incorporating cells lacking informative lineage data. Pseudotime analysis allows for the continuous nature of cell state transitions to be analysed, as opposed to relying on discrete experimental longitudinal sampling. This allows us to maximise the data that it is possible to capture experimentally by using this ensemble view to approximate for the trajectory taken by a single cell. Furthermore, this facilitates identification of genes underlying post-treatment Pro-to-Hep plasticity characterised above (which we refer to as ‘recovery plasticity’), defining a set of ‘recovery plasticity markers’.

We applied Slingshot (Street *et al*, 2018) analysis to assign HuH6 cells a pseudotime ordering representative of recovery plasticity (Figure 5A). Across three recovery pseudotime trajectories inferred by Slingshot, recovery plasticity markers were those that exhibited expression patterns correlating with pseudotime progression (Figure EV3A). We defined recovery plasticity markers operationally as genes whose expression was significantly upregulated (termed ‘recovery-up’) or downregulated (‘recovery-down’) along the Pro-to-Hep recovery pseudotime (Supplementary Data 2).

**Figure 5:**
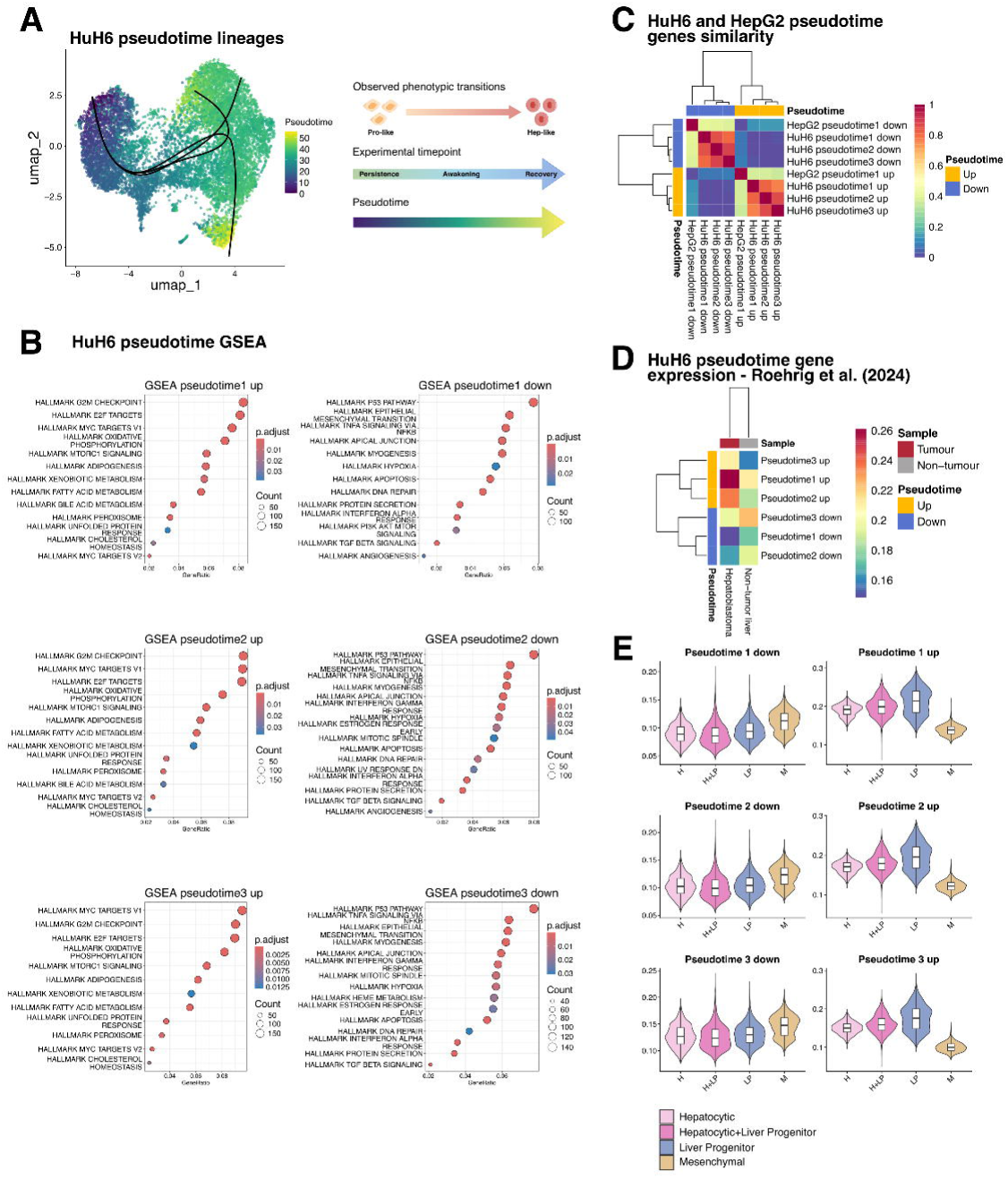
**(A)** scRNA-seq UMAP of 10,791 HuH6 cells that passed QC, coloured by computed shared pseudotime value. Black lines represent recovery lineages 1-3. Schematic depiction of the relationship between the observed phenotypic transitions, experimental phases and pseudotime. **(B)** GSEA plots for MSigDB Hallmark terms that are associated with the genes whose expression, either increasing (up) or decreasing (down), correlated with pseudotime progression. **(C)** Jaccard similarity between the pseudotime-derived genes identified in the HuH6 and HepG2 scRNA-seq data. Colour indicates the fraction of shared genes between each pair of sets, from 0 (no overlap) to 1 (identical). **(D)** Heatmap of the module scores for recovery plasticity markers in scRNA-seq data from six patient samples (Roehrig *et al*, 2024b). Expression in pseudobulked tumour and non-tumour annotated cells, where colour indicates mean expression of signature genes relative to a matched control set. **(E)** Expression of plasticity markers in cells, grouped by cell states defined in Roehrig *et al*. (2024): Hepatocytic (H), Hepatocytic+Liver Progenitor (H+LP), Liver Progenitor (LP), and Mesenchymal (M).

Gene set enrichment analysis (GSEA) showed that recovery-up genes were enriched for liver metabolic processes (e.g. *SCD*, *AFP*, *APOA1*) and MYC-/E2F-driven proliferation programs (e.g. *TOP2A*, *NCL*, *HMGB2*), while recovery-down genes were enriched for EMT (e.g. *CD44*, *SERPINE1*, *SAT1*) and TP53-stress responses, such as DNA repair and apoptosis (e.g. *MDM2*, *ATF3*, *SFN*; Figure 5B). The co-occurrence of TP53 target enrichment, wild-type in most HB tumours (Nagae *et al*, 2021; Eichenmüller *et al*, 2014), and stress response modules in recovery-down genes is consistent with cisplatin-induced, TP53-mediated arrest sustaining the persister state, while exit from persistence involved MYC/E2F reactivation, consistent with the release of TP53-mediated blockade.

To dissect how plasticity and proliferation programs change across the awakening-to-recovery period, we first examined their population-level kinetics. Within recovery-up genes, we identified two subsets – Hep-like identity genes (derived from HuH6 DGE analysis; see Methods and Supplementary Data 1) and cell cycle genes. The mean population-level expression across experimental phases revealed distinct kinetics: Hep-like identity markers collapsed sharply during persistence and recovered ahead of cell cycle re-engagement, with cell cycle markers recovering more gradually (Figure EV3B). This indicates that re-acquisition of Hep-like identity precedes full proliferation re-entry at the population level. To test whether plasticity and proliferation are coupled within individual cells, we computed per-cell module scores for two Hallmark gene sets: EMT as a marker of plasticity and G2M checkpoint as a marker of proliferation. Cells stratified along a plasticity-proliferation axis with strong inverse correlation at the single-cell level: cells occupied predominantly either an EMT-high/G2M-low or an EMT-low/G2M-high regime, with very few cells occupying both states simultaneously (Figure EV3C). Persister cells (Cisplatin condition) clustered in the EMT-high/G2M-low, whereas Recovery t_3_ cells clustered in the EMT-low/G2M-high regime, overlapping in their EMT-G2M coordinates with pre-treatment populations (POT, Untreated), consistent with phenotypic re-diversification that resembles a pre-treatment-like state. Cells from intermediate recovery timepoints (Recovery t_1_, Recovery t_2_) were distributed across the transition between both regimes, consistent with a progressive exit persistence. This single-cell anti-correlation indicates that plasticity (EMT) and proliferation (G2M) operate in near mutually exclusive regimes within individual cells: the persister phase is enriched with cells exhibiting a Pro-like highly plastic yet arrested phenotype that is exited through an initial cell state switch (Pro-to-Hep) followed by proliferation re-entry.

We applied the same pseudotime analysis pipeline to the HepG2 data to further interrogate the Pro-to-Hep dynamics identified by lineage-traced transcriptomes. Slingshot pseudotime inference identified an analogous Pro-to-Hep recovery trajectory (Figure 5C; Figure EV3D). HepG2 recovery-up genes were again enriched for MYC-/E2F-driven proliferation programs, while recovery-down genes were enriched for EMT (including TGF-β programs), DNA repair and apoptosis (Figure EV3E; Supplementary Data 3). The conservation of gene set associations within recovery plasticity markers derived from HuH6 and HepG2 models supports a shared transcriptional program underlying Pro-to-Hep plasticity-driven recovery in these HB cell line models.

To investigate the clinical relevance of recovery plasticity markers, we analysed their expression in a published scRNA-seq dataset of six HB patient samples (Roehrig *et al*, 2024a, 2024b). Within HB tumour cells, recovery-up signature scores were more elevated in tumour compared to non-tumour liver, whereas downregulated markers (recovery-down) were depleted in tumours and expressed in non-tumour liver (Figure 5D). Recovery-up signature scores were elevated across the hepatic-epithelial states – highest in the liver progenitor (LP) state and also elevated in the hepatocytic (H) state – and lowest in the mesenchymal (M) state, consistent with the combined proliferative and hepatocyte-identity content of the recovery-up program. Recovery-down scores showed the reciprocal pattern, being elevated in the mesenchymal (M) state (Figure 5E). This distribution indicates that the recovery plasticity signatures capture transcriptional programs relevant to multiple phenotypic states described in HB patient tumours; a direct demonstration of LP-to-H or M-to-LP transitions in patient tumours would, however, require large cohorts of longitudinal patient samples beyond the scope of this study.

### Cisplatin changes the phenotypic landscape and enables new transition routes coupled to plasticity and proliferation programs

To complement the pseudotime analysis with an orthogonal view of phenotypic dynamics, we applied dynamic modelling using MuTrans, a method that distinguishes transitioning cells from cells residing in stable attractor basins (Zhou *et al*, 2021). Unlike pseudotime inference, this dynamical modelling approach does not presume continuous cell state transitions; it instead models a multi-scale switch process between attractors via transition cells. In doing so, MuTrans visualises population dynamics as Waddington-like landscapes and computes, for each cell, a transition entropy, a measure of how confidently the cell can be assigned to a single attractor basin. Cells with high transition entropy sit near saddle points between basins and are interpreted as transitioning between states, while cells with low entropy occupy stable attractor basins, interpreted as ‘committed’ to a state. We applied MuTrans separately to pre-treatment (POT, Untreated) and post-treatment (Cisplatin, Recovery timepoints) HuH6 and HepG2 datasets, annotating attractor basins by the developmentally defined states previously applied (Figure EV4A; also see Figure 1). Informed by the developmental roots of these states, we defined a canonical topology as one in which less stable Pro-like states occupy shallow basins that flow into stable deeper Hep-like attractors, paralleling Waddington’s 1957 model of differentiation (Waddington, 1957), and asked whether cisplatin treatment produces deviations from this topology.

In pre-treatment HuH6 populations, the inferred landscape resembled canonical differentiation with Pro-like states occupying shallow basins that flowed into deeper Hep-like attractors, recapitulating normal differentiation landscapes (Figure 6A). Post-treatment landscapes were restructured: differences between attractor basin depths decreased and slopes flattened, favouring persistence in Pro-like states and reducing the bias toward the Pro-to-Hep ‘valleys of terminal differentiation’ seen pre-treatment (Figure 6B; Appendix Table 5). Post-treatment cells were no longer biased towards Hep attractors but distributed across multiple Hep- and Pro-like basins with comparable depth.

**Figure 6:**
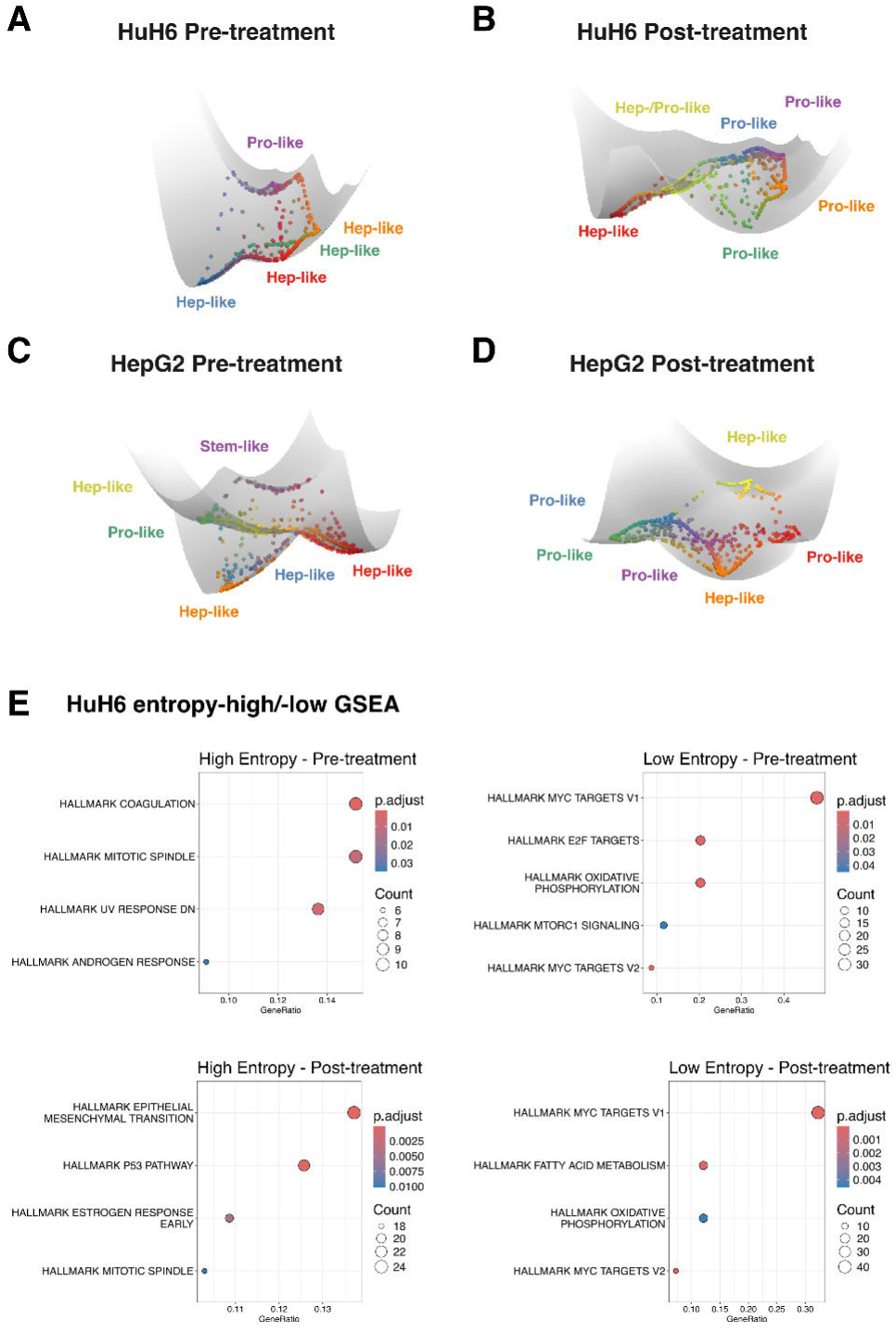
**(A)** Dynamical manifold of HuH6 cells pre-treatment (POT, Untreated) coloured by MuTrans attractor. Labels indicate the predominate cell state of the attractor wells as annotated by comparison with cell state identify previously derived from scRNA-seq analysis. **(B)** Dynamical manifold of HuH6 cells post-treatment (Cisplatin, Recovery t_1_, Recovery t_2_, Recovery t_3_) coloured by MuTrans attractor. **(C)** Dynamical manifold of HepG2 cells pre-treatment (POT, Untreated) coloured by MuTrans attractor. **(D)** Dynamical manifold of HepG2 cells post-treatment (Cisplatin, Recovery t_1_, Recovery t_2_) coloured by MuTrans attractor. **(E)** GSEA plots for MSigDB Hallmark terms that are associated with the genes differentially expressed in high or low entropy HuH6 cells in the pre- or post-treatment datasets.

HepG2 cells displayed the same landscape restructuring (Figure 6C, D; Appendix Table 6). Pre-treatment HepG2 landscapes recapitulated canonical differentiation topology, with the rare Stem-like state occupying the highest basin and feeding into Pro-like and downstream Hep-like attractors. Post-treatment HepG2 landscapes showed the same slope flattening and basin depth reduction observed in HuH6. The conservation across both cell line models supports that landscape restructuring is a general feature of HB cisplatin response.

To identify transcriptional programs accompanying cell state transitions, we compared transitioning cells (high entropy) versus committed cells (low entropy) by performing differential gene expression and GSEA (Figure EV4B). This revealed an enrichment for MYC- and E2F-target programs in committed cells, while EMT-related pathways were enriched in transitioning cells post-treatment (Figure 6E; Supplementary Data 4). These same gene sets were identified in pseudotime trajectory analysis (see Figure 5B) – MYC-/E2F-driven proliferation in recovery-up genes, EMT programs in recovery-down – providing orthogonal confirmation that the plasticity-proliferation axis underlying recovery is detectable by two methodologically distinct approaches, arguing against a method-specific artefact.

### Integrated regulatory and chromatin analysis identifies convergence on *BIRC5* as a candidate node of the proliferation-plasticity axes

To identify candidate transcriptional regulators of recovery plasticity, we performed motif enrichment analysis on recovery-up genes in HuH6, asking which transcription factor (TF) binding motifs were overrepresented in their proximal regulatory regions (Figure 7A). We identified 24 TF motifs enriched across all three recovery-up gene sets, derived from the three pseudotime lineages (Figure EV5A; Supplementary Data 5). The same 24 motifs were also enriched in HepG2 recovery-up genes, supporting their conservation across cell line models.

**Figure 7:**
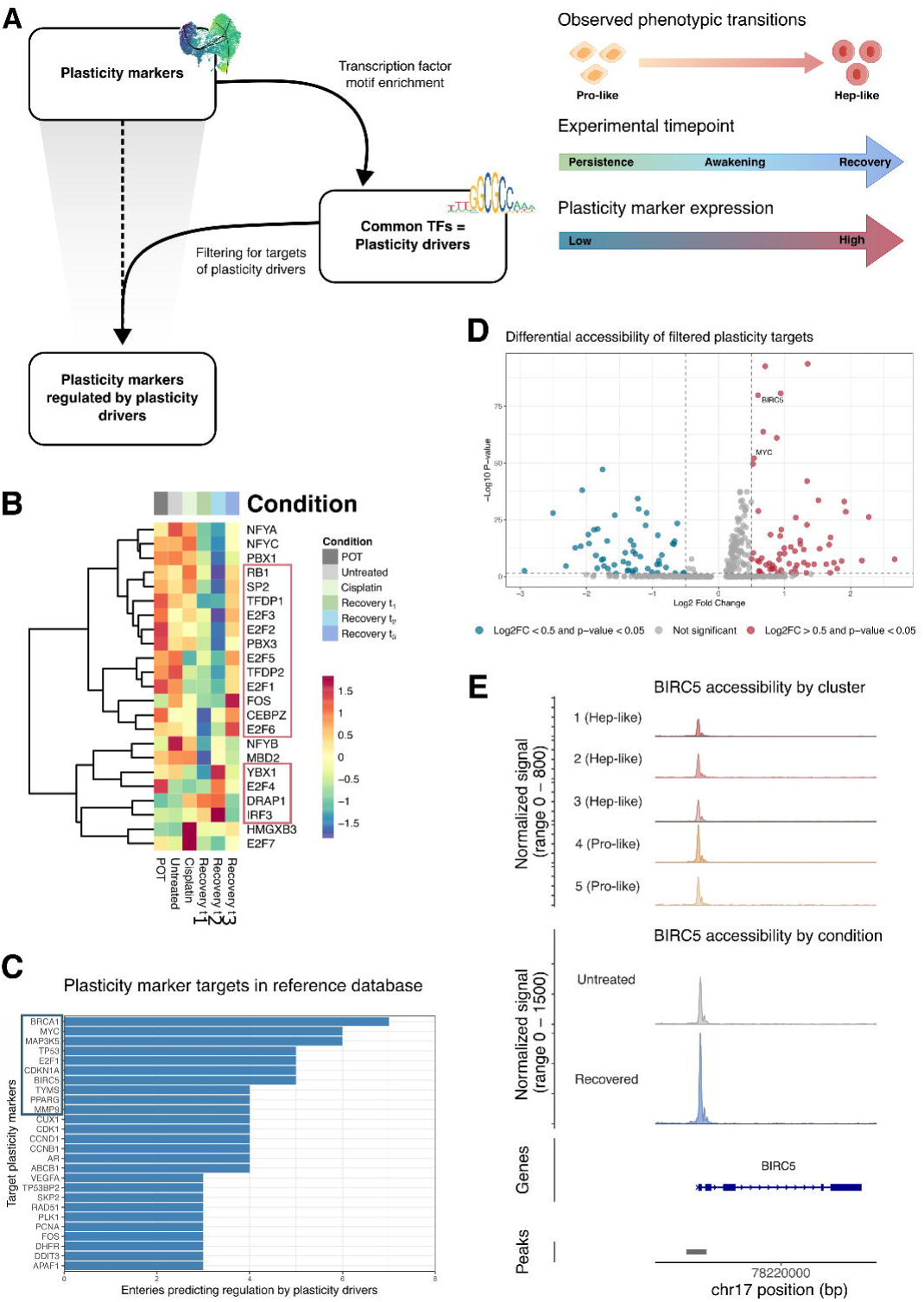
**(A)** Strategy to identify plasticity drivers and targets. (1) Transcription factor motif enrichment on plasticity markers (recovery-up genes). (2) Identification of plasticity drivers – candidate transcription factors identified across recovery-up gene lists. (3) Filtering of plasticity markers (recovery-up genes) by known targets of plasticity drivers. **(B)** Expression of the 23 shared plasticity drivers (expression data for *FOXI1* was unavailable) at each experimental timepoint in the HuH6 scRNA-seq data. **(C)** Number of entries for each target from querying a reference database of transcription factor regulatory relationships (Han *et al*, 2018) for the plasticity drivers, with the top 10 hits highlighted. **(D)** Differential accessibility analysis between recovered and untreated samples of plasticity targets, with thresholds of log_2_ fold change > 0.5 and adjusted *p*-value < 0.05. Labelled points refer to genes that were in the top 10 hits from our reference database query. **(E)** Averaged frequency of sequenced DNA fragments associated with *BIRC5* (top) grouped by snRNA-derived phenotype, (bottom) grouped by experimental condition.

We reasoned that genuine drivers of recovery must be expressed before or concurrent with the re-expression of their target genes during recovery. Recovery-up genes were expressed at comparable levels in untreated and recovered populations but reduced during persistence and awakening (Figure EV5B): 16/24 TFs showed peak expression at Recovery t_2_ or t_3_, compatible with regulating their downstream recovery-up targets (Figure 7B). To go beyond motif enrichment to literature-validated regulation, we cross-referenced the 16 candidate TFs against a manually curated database of human transcriptional regulatory networks (Han *et al*, 2018). For each TF, we identified the subset of recovery-up genes it had previously been shown to regulate, yielding a high confidence set of TF-target links within the recovery-up program. The top hits were recovery-up genes belonging to MYC/E2F and TP53 pathways (Figure 7C), linking the candidate plasticity driver TFs to the plasticity-proliferation axis and motivating focused analysis of these targets in the chromatin analysis and functional experiments that follow.

To examine whether these regulatory dynamics leave an accessibility signature at the chromatin level, we generated single-nuclei ATAC sequencing (snATAC-seq) data from HuH6 cells at the Untreated and Recovery t_3_ timepoints as part of a multiomic profile, allowing direct comparison of chromatin accessibility before and after the recovery cycle. We first asked how chromatin accessibility at canonical HB marker loci (*AFP*, *CTNNB1*, *HNF4A*, *LEF1*, *TCF4*) differed between untreated and recovered cells (Figure EV5C). Most HB marker loci showed reduced accessibility in recovered cells, indicating chromatin-level loss of pre-treatment HB identity at these regions. Two notable exceptions were *HNF4A* and *LEF1*, key downstream effectors of Wnt/β-catenin, the principal oncogenic driver in HB and a regulator of hepatocytic differentiation (Kluiver *et al*, 2024), which retained accessibility at their loci in recovered cells. This selective retention is consistent with Wnt/β-catenin signalling remaining active during recovery and supports its role in regulating the transcriptional dynamics of plasticity-driven phenotypic transitions.

In contrast to canonical HB markers, recovery-up genes regulated by the 16 candidate plasticity driver TFs showed increased chromatin accessibility in recovered cells (Figure 7D). Targets of ten TFs (E2F1, E2F3, E2F4, E2F6, FOS, IRF3, RB1, SP2, TFDP1, YBX1) were significantly more accessible in the Recovery t_3_ compared to the Untreated sample (Supplementary Data 6). This is particularly striking because the global transcriptome of Recovery t_3_ cells largely reverts to a pre-treatment state (see Figure EV1A). The chromatin and transcript level therefore decouple at recovery-up gene loci, with the retained accessibility being consistent with an epigenetic memory of the cisplatin recovery cycle. Among the top recovery-up targets from our prioritisation, *MYC*- and *BIRC5*-associated loci showed this gained and retained accessibility pattern, raising the possibility that these loci form a plasticity-first adaptation program – primed at the chromatin level to enable faster phenotypic reemergence on subsequent therapy exposure.

*BIRC5* emerges from this prioritisation as a particularly interesting candidate. *BIRC5* is a β-catenin target with established roles in cell survival and proliferation (Kelly *et al*, 2011), and is a marker of the aggressive C2 subtype of HB (Cairo *et al*, 2008); while *BIRC5* has previously been proposed as a therapeutic target in HB (Yu *et al*, 2015), our data suggests a distinct mechanistic framing – *BIRC5* as a node coupling plasticity to proliferation re-entry, with epigenetic priming of its locus during recovery. Specifically, aggregated chromatin accessibility profiles showed that the *BIRC5* locus was more accessible in Pro-like cells, the phenotype that survives cisplatin – priming the locus ahead of its later transcription in proliferative, Hep-committed cells – and this accessibility was retained in recovered cells relative to untreated controls (Figure 7E). This occurred despite *BIRC5* transcript levels being comparable between the Untreated and Recovery t_3_ cells (Figure EV5D), indicating the *BIRC5* locus is epigenetically primed rather than constitutively transcriptionally active.

Together, these findings suggest a regulatory model linking cisplatin response to plasticity-driven recovery via the E2F-*BIRC5* axis. Cisplatin-induced p53 activation represses E2F activity (Benson *et al*, 2014; Fischer *et al*, 2015), through p21- and p16-mediated CDK inhibition, Rb sequestration, and DREAM-complex repression of E2F targets, including *BIRC5*. As DNA damage is repaired and p53 activity declines during recovery, E2Fs are released, *BIRC5* transcription becomes competent and proliferation re-entry, in coordination with phenotypic transitions toward Hep-like attractor states, ensues with sustained Wnt/β-catenin activity (see above) supporting hepatocytic commitment alongside E2F-driven *BIRC5* re-expression. The retained chromatin accessibility at recovery-up loci, including *BIRC5*, indicates that this re-activation is poised to occur on subsequent therapy exposure – defining the possibility of a plasticity-first adaptation framework in which epigenetic priming of axis genes accumulates across treatment cycles. In the context of the plasticity-proliferation axis suggested by our pseudotime (see Figure 5) and landscape analysis (see Figure 6), this combined regulatory and multiomics analysis positions *BIRC5* as an axis driver, which we investigate functionally below by both genetic and pharmacological perturbation.

### *BIRC5* maintains hepatocytic-like phenotypes and represents a cisplatin-persister vulnerability

*BIRC5*, the gene encoding Survivin, is part of the inhibitor of apoptosis (IAP) protein family, is an actionable target (Li *et al*, 2019), is expressed highly across tumour types (O’Connor *et al*, 2000) and is a marker of the C2-subtype in HB (Cairo *et al*, 2008). To confirm its clinical relevance, we examined *BIRC5* expression in HB patient material and its association with prognostic features. In the Carrillo-Reixach *et al*. (2020) discovery cohort samples (32 tumour and 32 paired non-tumour, NT) (Carrillo-Reixach *et al*, 2020a, 2020b), *BIRC5* expression was significantly higher in tumour than NT liver (Figure 8A). High *BIRC5* expression in post-treatment surgical specimens was associated with the aggressive C2 subtype. Elevated *BIRC5* was also associated with other molecular features of poor prognosis, such as strong expression of the 14q32 gene signature, the Epi-CB epigenetic subtype – characterised by global DNA hypomethylation and CpG island hypermethylation (Carrillo-Reixach *et al*, 2020b) – and the high-risk molecular risk subclass (MRS-3) (Table 2). Stratifying patients by median *BIRC5* expression revealed that high-*BIRC5* patients had significantly lower overall survival (*p* = 0.0009; Figure 8B) and a higher incidence of recurrence or death. Together, these findings establish *BIRC5* as both a marker of adaptive post-treatment potential and a clinically relevant predictor of poor prognosis in HB.

**Figure 8:**
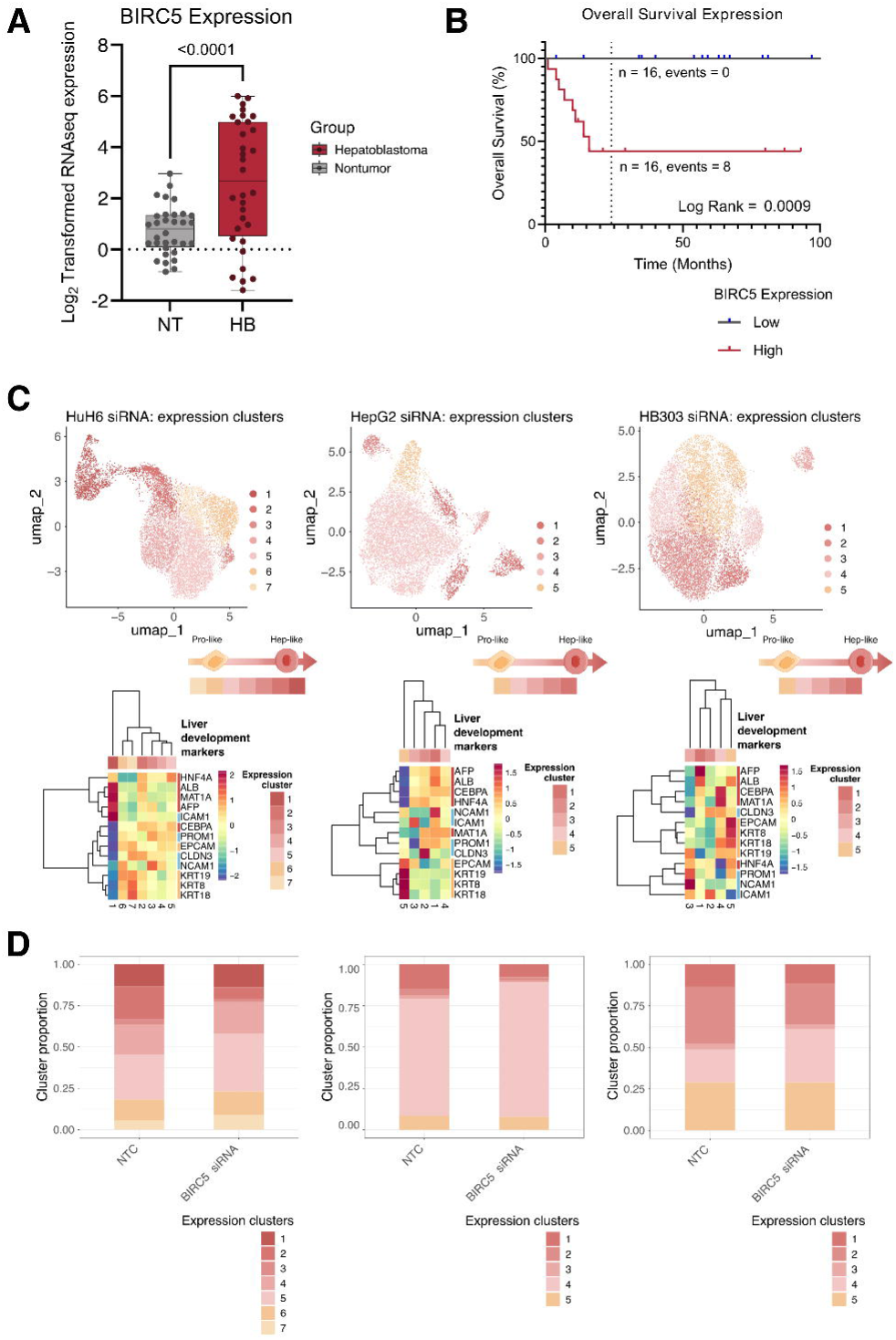
**(A)** Expression of plasticity target *BIRC5* in RNA sequencing (RNA-seq) data of 32 HB tumours and 32 paired non-tumour (NT) samples (Carrillo-Reixach *et al*, 2020b). *p*-values were calculated using paired Wilcoxon test and considered significant if *p* < 0.05. **(B)** Overall survival (OS) Kaplan-Meier plot of patients categorised according to *BIRC5* expression, based on median tumour expression. **(C)** scRNA-seq UMAPs of three cell lines – (left-to-right) HuH6, HepG2, HB303 – that were transfected with either a non-targeting control (NTC) or *BIRC5* siRNA, coloured by expression cluster. The number of cells that passed QC and were plotted in each UMAP are as follows: HuH6 10,726; HepG2 8,197; HB303 11,588. **(D)** Proportion of each expression cluster in each cell line population – (left to right) HuH6, HepG2, HB303 – split by experimental condition (transfection with NTC or *BIRC5* siRNA).

To test whether *BIRC5* functions within the plasticity-proliferation axis, we examined how *BIRC5* transcript expression is distributed across cells relative to EMT and G2M signature expression (see Figure EV3C). Across all cells, *BIRC5* expression was significantly positively correlated with Hallmark G2M module scores (*r_s_* = 0.61; *p* < 0.05) and negatively correlated with Hallmark EMT module scores (*r_s_* = −0.34; *p* < 0.05; Figure EV6A). *BIRC5*-high cells localised to an EMT-low/G2M-high regime. This places *BIRC5* expression on the proliferation arm of the axis at single-cell resolution, consistent with its role as an E2F-target gene and indicating that *BIRC5* expression is coupled to the proliferative, Hep-committed phenotype and is lowest in the persister Pro-like state. This predicts a requirement for *BIRC5* in maintaining Hep-like states, which we tested directly below.

To functionally test whether *BIRC5* is required to maintain committed Hep-like states, we performed siRNA knockdown of *BIRC5* in three preclinical models – HuH6, HepG2, and the PDX-derived HB303 line – and assessed the phenotypic composition of the resulting populations by scRNA-seq (Figure EV6B; Appendix Table 7). Knockdown efficiency was confirmed by western blot in all three models (Figure EV6C). Biological duplicates underwent scRNA-seq and clusters were annotated, as previously outlined, using relevant HB signatures (Carrillo-Reixach *et al*, 2020b; Roehrig *et al*, 2024b; Cairo *et al*, 2008), and liver development signatures (Wesley *et al*, 2022) and markers (Lemaigre, 2009; Si-Tayeb *et al*, 2010; Schmelzer *et al*, 2007; So *et al*, 2020; Ramani & Lu, 2017) from literature (Figure 8C, Figure EV6D). In HuH6, *BIRC5* knockdown produced a shift towards less differentiated, more progenitor-like phenotypes, characterised by increased expression of hepatoblast markers (*KRT8*, *KRT18* and *KRT19*) and co-expression of Wesley hepatocyte and cholangiocyte signatures. In HepG2 and HB303 data, the effects were more subtle: a cluster (4 for both) expressing the Wesley hepatocyte signature with reduced *ALB* and *AFP* and elevated hepatoblast marker (*EPCAM* and *KRT*s) expression dominated after *BIRC5* knockdown, consistent with a partial Hep-to-Pro shift. Together this data indicates that *BIRC5* supports Hep-like phenotype maintenance in steady-state conditions, with model-dependent magnitude.

To test whether pharmacological *BIRC5* inhibition could exploit the post-treatment vulnerability suggested by our regulatory and functional analyses, we used YM155 (sepantronium bromide) (Nakahara *et al*, 2007), a small molecule that suppresses *BIRC5* transcription, reportedly by interfering with SP1-mediated promoter activity at the *BIRC5* locus. YM155 is the first of its class and the most extensively characterised *BIRC5*-targeting compound, having progressed to clinical testing in adult cancers (Clemens *et al*, 2015; Satoh *et al*, 2009; Tolcher *et al*, 2008).

Based on the regulatory and chromatin findings (see Figure 7), we hypothesised that *BIRC5* inhibition may exert a stronger effect in cisplatin-induced persister cells than in untreated cells, regardless of single-agent sensitivity. To test this, we compared YM155 viability effects in the HuH6, HepG2, and HB303 cell lines that were either treatment-naive or pre-exposed to cisplatin to generate a persister-enriched population. To establish dose ranges for combination experiments, we first performed drug-response assays (Figure EV6E). These demonstrated that their sensitivities to YM155 differed substantially across models: HuH6 showed dose-dependent loss of viability at the tested YM155 concentrations, while HepG2 and HB303 were comparably insensitive across the same ranges. This variable single-agent activity parallels the variable effect of *BIRC5* siRNA knockdown across cell lines models (see Figure 8D).

Cisplatin-induced persister cells were sensitive to YM155 in all three cell lines, with effects that exceeded those of YM155 as a single agent (Figure 9A). In HuH6, YM155 alone approximately accounted for 96% cell death, while addition of YM155 to cisplatin-tolerant persister cells produced 99% cell death. In HepG2 and HB303, models insensitive to YM155 alone, addition of YM155 to cisplatin pre-treated cells produced over 30% additional loss of viability over cisplatin alone. This persister-specific vulnerability indicates that the combinatorial effect is not simply explained by baseline drug sensitivity. Recovery from sequential treatment compared to cisplatin alone was also observed to be impeded across all three cell lines (Figure 9B; Supplementary Data 7).

**Figure 9:**
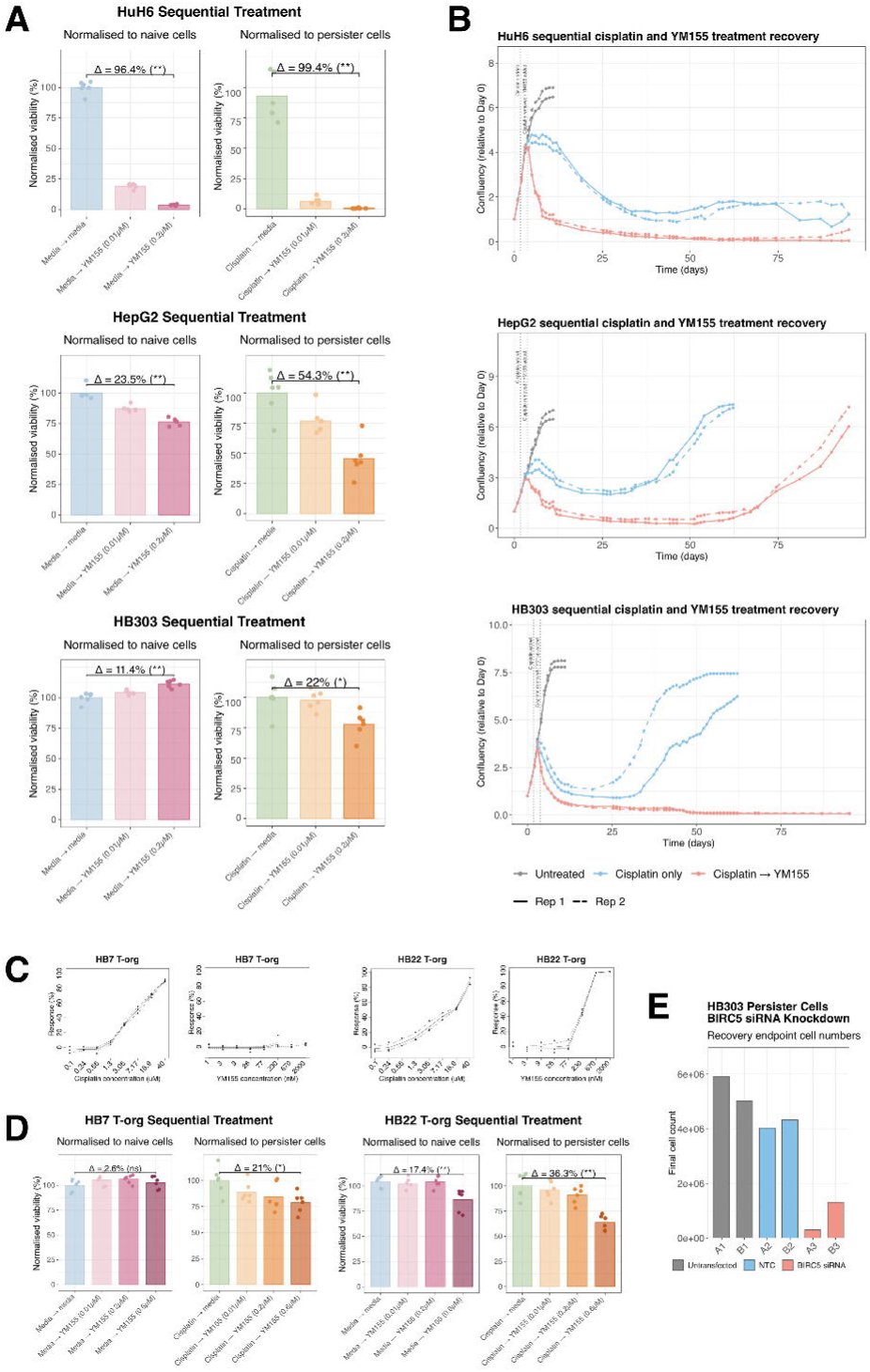
**(A)** Sequential treatment in (top-to-bottom) HuH6, HepG2, and HB303: first with media or cisplatin (IC50), then with media or YM155 at two concentrations (0.01µM, 0.2µM). The media first (cisplatin naive) conditions were normalised to media + media control group; the cisplatin first (cisplatin persister) conditions were normalised to cisplatin + media control group. The greatest percentage difference comparisons between the relevant control group (media/cisplatin + media) and the corresponding media/cisplatin + YM155 (0.2µM) are annotated as Δ. The *t*-test results for these differences are also annotated as the following: ns = non-significant, * = *p* < 0.05, ** = *p* < 0.01. **(B)** Confluency of (top-to-bottom) HuH6, HepG2, and HB303 cultures under three conditions: (grey) untreated, (blue) cisplatin only (IC50, 48 hours), (red) cisplatin (IC50, 48 hours) followed by YM155 (0.01µM for HuH6 and HepG2, 0.2µM for HB303). **(C)** Drug-response curves in HB7 T-org (left) and HB22 T-org (right) for cisplatin (concentration ranges from 0.01 to 40µM) and YM155 (concentration ranges from 1 to 2000nM). In all cases, the first point (left) is the vehicle. **(D)** Sequential treatment in HB7 T-org (left) and HB22 T-org (right): first with media or cisplatin (25µM), then with media or YM155 at three concentrations (0.01µM, 0.2µM, 0.6µM). The media first (cisplatin naive) conditions were normalised to media + media control group; the cisplatin first (cisplatin persister) conditions were normalised to cisplatin + media control group. The greatest percentage difference comparisons between the relevant control group (media/cisplatin + media) and the corresponding media/cisplatin + YM155 (0.6µM) are annotated as Δ. The *t*-test results for these differences are also annotated as the following: ns = non-significant, * = *p* < 0.05, ** = *p* < 0.01. **(E)** Final cell count at the recovery endpoint (35 days post-treatment) of cisplatin-treated (IC50) HB303 cultures left to recover, either untransfected or transfected with a non-targeting control (NTC) or *BIRC5* siRNA (re-transfected every three weeks to maintain knockdown over the recovery period). Two biological replicates (A, B) are shown per condition.

To support these observations, we carried out an analogous combination experiment in two patient-derived organoid (PDO) cultures (Clavería-Cabello *et al*, 2023). Drug-response curves for cisplatin and YM155 on PDO cultures (HB7 T-org and HB22 T-org) showed the same model-dependent sensitivities to YM155 killing (Figure 9C). While both PDO-derived models were similarly responsive to cisplatin, their sensitivities to YM155 differed substantially with HB22 T-org (like HuH6) showing dose-dependent loss of viability, while HB7 T-org (like HepG2 and HB303) was insensitive across the same range.

In PDO lines, we further observed evidence that cisplatin-induced persister cells were sensitive to YM155 in both PDO lines (Figure 9D). In HB7 T-org (completely insensitive to YM155 alone) addition of YM155 to cisplatin pre-treated cells produced more than 20% additional loss of viability over cisplatin alone. In HB22 T-org, YM155 alone approximately accounted for 17% cell death, while addition of YM155 to cisplatin-tolerant persister cells produced 36% cell death, greater than two-fold enhancement over single-agent activity. The same persister-specific vulnerability observed in cell lines was observed in PDO lines, regardless of whether the model was sensitive (HB22 T-org) or insensitive (HB7 T-org) to single-agent YM155, further supporting that the sequential effect is not dictated by model-dependent sensitivity to YM155 as a single-agent.

To test whether the persister-context vulnerability reflects an on-target *BIRC5* dependency rather than an uptake- or off-target-dependent effect of YM155, we downregulated *BIRC5* by siRNA in cisplatin-treated persister cells in HB303, a clean background as YM155 had no activity in untreated HB303 cells over the concentrations tested. Downregulation of *BIRC5* markedly reduced recovery-endpoint (35 days post-treatment) cell numbers relative to the NTC (Figure 9E). Together, the genetic and pharmacological data attribute the persister-context vulnerability specifically to *BIRC5*, while indicating that its therapeutic exploitation will depend on effective target engagement.

In summary, this data supports a model in which *BIRC5* maintains the proliferation-coupled Hep-like phenotype in steady-state HB cells and constitutes a context-dependent vulnerability in cisplatin-persister cells. The dissociation between single-agent YM155 activity and persister-context efficacy suggests that *BIRC5* inhibition exploits a dependency that emerges during cisplatin recovery, consistent with the persistent epigenetic priming at the *BIRC5* locus (see Figure 7). These findings position *BIRC5* as a candidate target for sequential, combinatorial intervention in HB, with the potential to selectively target the persister population that fuels plasticity-led recovery and may underlie relapse.

## Discussion

Outcomes for patients with chemo-resistant HB remain poor, with limited therapeutic options. A more fundamental understanding of the evolutionary processes underlying treatment adaptation is needed to guide new strategies. The intrinsic heterogeneity of cancer has long been proposed as a driver of resistance (Hanahan & Weinberg, 2011; Greaves, 2015), but whether this heterogeneity reflects pre-existing states or therapy-induced adaptation has remained unresolved, largely due to technological limitations in resolving clonal and temporal dynamics. This distinction has direct clinical implications: should we focus on eradicating pre-existing resistant clones, or on steering tumours towards extinction through adaptive therapies? In HB, specifically, the low mutational burden and high developmental phenotypic heterogeneity make this distinction especially consequential, as they implicate the role of non-genetic plasticity-driven mechanisms in treatment failure and relapse.

By combining expressible DNA barcoding with longitudinal single-cell transcriptomics across the persistence-awakening-recovery course, we dissociate phenotypic from clonal dynamics during cisplatin response in HB. Our central finding is that the two operate on different scales: cisplatin imposes a strong phenotypic bottleneck, selecting for progenitor-like cells across all clones, but does not produce a strong clonal sweep. Different subsets of persister clones resume proliferation in different replicates with a common recovery window, with only modest reproducible fitness biases among shared clones. Exit from persistence therefore occurs through asynchronous, partly fitness-biased clonal re-expansion, coupled with plasticity-led phenotypic re-diversification toward the pre-treatment state. This framework of phenotypic selection at the state level, asynchronous and partly stochastic selection at the clonal level refines models of treatment resistance and clarifies the actionable target population in chemoresistant HB: the small persister pool that fuels recovery, not necessarily a pre-existing genetically resistant clone.

Phenotypic plasticity has been increasingly recognised as a driver of resistance (Da Silva-Diz *et al*, 2018; Qin *et al*, 2020). In paediatric cancers, phenotypic plasticity is of particular interest due to the developmental origin of childhood tumours and the intrinsic plasticity of their cells of origin (West-Eberhard, 2003; Hanahan, 2022). The pre-existing intra-clonal phenotypic heterogeneity observed in our models (present at the Untreated timepoint without selective pressure) supports a model in which therapy acts on pre-existing transcriptionally distinct cell states rather than inducing them, consistent with phenotypic noise serving as a substrate for selection (Whiting *et al*, 2024).

Our longitudinal patient case study, while limited to two cases, demonstrates that the less-to-more differentiated-like direction of post-treatment dynamics is conserved across HB subtypes despite their distinct molecular and histological backgrounds: P1 samples, from a C1-like tumour, show movement along a hepatocytic-to-hepatoblast axis, whilst P2 samples, from a C2-like tumour, additionally involves a less-differentiated, bipotent progenitor-like state. This subtype-dependent scope of plasticity refines previous descriptions of HB phenotypic heterogeneity (Cairo *et al*, 2008; Carrillo-Reixach *et al*, 2020b; Hirsch *et al*, 2021; Roehrig *et al*, 2024b; Song *et al*, 2022; Hooks *et al*, 2018; Sekiguchi *et al*, 2020; Sumazin *et al*, 2022; Nagae *et al*, 2021; Chen *et al*, 2025) by highlighting how subtype context constrains the range over which post-treatment plasticity operates.

At single-cell resolution, we found plasticity (EMT) and proliferation (G2M) module scores were anti-correlated, with cells occupying near mutually exclusive regimes: persister cells were plasticity-high/proliferation-low, while recovered cells were plasticity-low/proliferation-high. The persister state thus represents a plastic-arrested phase that must be exited before proliferation can re-engage. This suggests the existence of a temporal ordering in which plasticity precedes proliferation re-entry rather than emerging concurrently. Consistent with this, within the recovery-up gene set, hepatocytic-like identity markers fell dramatically and re-engaged ahead of cell-cycle markers at the population level, indicating that re-acquisition of hepatocytic-like identity precedes proliferation re-entry. These observations refine previous reports that chemotherapy-induced stem-like HB cells can switch between subtypes to promote repopulation; they support at single-cell resolution that a plasticity-proliferation axis underlies this phenomenon (Lee-Theilen *et al*, 2022).

Mechanistically, integrated regulatory and chromatin analyses identified E2F transcription factors as candidate drivers of recovery-up programs, with *BIRC5* emerging as a key E2F-target node linking plasticity to proliferation re-entry. It is known that cisplatin-induced p53 activation results in p21- and p16-mediated CDK inhibition (Benson *et al*, 2014; Chen *et al*, 1996; McConnell *et al*, 1999), Rb sequestration, and DREAM-complex repression of E2F target genes (Fischer *et al*, 2015), such as *BIRC5*. Subsequent repair of DNA damage can result in p53 activity decline during recovery. This can release E2Fs, facilitating *BIRC5* transcription. Positive feedback loops in which E2F1 promotes CDK expression further amplify this reactivation, allowing cells to escape p53-dependent arrest (DeGregori *et al*, 1995; Zetterberg *et al*, 1995), restart proliferation in coordination with committing to hepatocytic states. Single-cell expression analysis supports this regulatory model: *BIRC5* expression was positively correlated with proliferation module scores and negatively correlated with EMT scores, localising *BIRC5*-high cells specifically to the proliferation arm of the plasticity–proliferation axis. siRNA-mediated knockdown of *BIRC5* in untreated conditions shifted populations toward less-differentiated, progenitor-like phenotypes, providing causal evidence that *BIRC5* contributes to maintaining the proliferating hepatocytic-like state, with model-dependent magnitude.

Beyond transcriptional dynamics, multiomic profiling revealed that chromatin accessibility was maintained at the *BIRC5* locus after recovery, despite the global transcriptome largely reverting to a pre-treatment state. This decoupling of chromatin accessibility from transcript level indicates an epigenetic memory of the cisplatin–recovery cycle at these loci. We propose that this retained accessibility may constitute a plasticity-first adaptation program, in which repeated cisplatin exposure progressively primes recovery-up loci, including *BIRC5*, for faster transcriptional re-engagement on subsequent therapy exposure. Over multiple treatment cycles, such accumulating epigenetic priming could underpin the emergence of clinically resistant disease, consistent with the observed association between high *BIRC5* expression and poor patient survival, and with broader reports linking *BIRC5* to chemoresistance across cancer types, including adult liver cancer (Liu *et al*, 2006; Li *et al*, 2012; Fäldt Beding *et al*, 2022; Xu *et al*, 2021).

Functionally, *BIRC5* inhibition using the small-molecule YM155 produced a persister-specific vulnerability in patient-derived organoid models, most strikingly in HB7 T-org, where YM155 was inactive as a monotherapy yet produced substantial additional loss of viability when added to cisplatin-pretreated persister cells. This dissociation between single-agent activity and persister-context efficacy suggests that *BIRC5* inhibition exploits a dependency that emerges specifically after cisplatin treatment, consistent with the persistent epigenetic priming of the *BIRC5* locus. Genetic perturbation of *BIRC5* in cisplatin-treated HB303 persisters, a model refractory to YM155 as a single agent, reproduced this suppression of recovery, attributing the persister-context vulnerability specifically to *BIRC5* rather than to YM155’s off-target or delivery-dependent effects. While YM155 has known limitations as a tool compound, including dependence on the SLC35F2 transporter for cellular uptake (Winter *et al*, 2014) and reported off-target effects at higher concentrations (Glaros *et al*, 2012; Chang *et al*, 2015), the convergent evidence from the genetic (siRNA) and pharmacological (cisplatin + YM155) experiments supports *BIRC5* as a candidate combinatorial target for the minimal residual disease window in HB.

Several limitations should be noted: our patient analysis is limited to two longitudinal cases, of which sampling is of small portions of the tumours. Given the well-documented spatial heterogeneity of HB, one sample may not capture the full regional diversity of the tumour, and the differences observed between T1 and T2 could therefore reflect, in part, sampling of distinct tumour regions rather than temporal evolution alone. Additionally, one P2 sample is a lymph-node metastasis that may contribute biopsy-site effects alongside genuine post-treatment dynamics; larger longitudinal cohorts will be needed to establish whether these dynamics generalise across HB clinical contexts. We therefore interpret the patient data as illustrating a direction of post-treatment change that is concordant across two molecular subtypes and consistent with our controlled preclinical experiments, rather than as definitive per-patient trajectories. Our snapshot-based clonal sampling cannot resolve whether all persister clones retain latent proliferation potential given sufficient time, or whether a competence-determining process operates within the persister pool. The cell line models used, while powerful for the clonal lineage tracing analyses central to our study, occupy a C2-like, LP-like region of HB state-space and do not capture the full subtype heterogeneity of patient tumours, a limitation that we offset by case-study patient validation, but which warrants further work in HB models that capture C1-context dynamics.

Overall, our study provides a unified clonal, transcriptomic, and epigenetic view of plasticity-led adaptation in HB, demonstrating that recovery occurs through phenotypic selection of progenitor-like reservoirs, asynchronous clonal awakening, and epigenetically primed transcriptional regulation. Targeting plasticity-driven recovery in the minimal residual disease window, for example by combining differentiation-based therapies with inhibitors of proliferation-coupled axis components such as *BIRC5*, may provide a rational route to preventing relapse in HB and in other cancers where plasticity precedes resistance.

## Methods

### Cell culture

HuH6 (provided by Roland Kappler, Ludwig-Maximilians-Universität, Germany) and HepG2 (obtained from ECACC, Salisbury, UK) cell lines were cultured in Dulbecco’s Modified Eagle Medium (DMEM; 41965062, Gibco Carlsbad, CA, USA) supplemented with 10% foetal bovine serum (FBS; 10270106, Gibco), and 1% penicillin streptomycin (pen-strep; P4333, Sigma-Aldrich, St. Louis, MO, USA). HB303 (provided by Emilie Indersie, XenTech, France) were cultured in Advanced DMEM (12634010, Gibco) supplemented with 10% FBS, 1% pen-strep, 1% L-glutamine (G7513, Sigma-Aldrich), and 0.2% ROCK inhibitor (S1049, Stratech, Cambridge, UK). All cells were incubated at 37°C and 5% CO_2_. HB7 T-org and HB22 T-org PDO-derived cultures were cultured in BEC media: Advanced DMEM/F12 (12634010, Gibco) supplemented with 1% HEPES (15630056, Gibco), 1% GlutaMax™ (35050038, Gibco), 1% pen-strep (L0022, Biowest, Nuaillé, France), 1× N-2 supplement (11520536, Gibco), 1× B-27 supplement minus vitamin A (12587010, Gibco), 500ng/mL recombinant human (rh) RSPO1 (78213, STEMCELL Technologies, Vancouver, BC, Canada), 25ng/mL rhHGF (Z03229, GenScript, Nanjing, Jiangsu, China), 100ng/mL rhFGF10 (Z03314, GenScript), 50ng/mL rhEGF (Z00333, GenScript), 125ng/mL rhNoggin (120-10C, PreproTech, Cranbury, NJ, USA), 1.25mM *N*-acetylcysteine (A9165, Sigma-Aldrich), 10nM human [Leu^15^]-gastrin I (G9145, Sigma-Aldrich), 10mM nicotinamide (N0636, Sigma-Aldrich) and 5μM A 83-01 (1421, Tocris Bioscience, Bristol, UK), 0.5μM CHIR99021 (1386, Tocris Bioscience) and 10μM forskolin (2264, Tocris Bioscience).

### Lentiviral DNA barcoding

Transduction of founder HuH6 and HepG2 cells with the CloneTracker XP™ 1M Barcode-3’ Library with RFP-Puro (BCXP1M3RP-1S-P, Cellecta, Mountain View, CA, USA) was carried out as outlined in the CloneTracker XP™ lentiviral barcode libraries user manual. Optimisation to achieve a multiplicity of infection (MOI) of 0.1 was carried out. Transduced cells underwent fluorescence-activated cell sorting to isolate RFP-positive, barcoded cells. An initial population of 1.26 × 10^5^ barcoded HuH6 cells were expanded (×233) to form the POT, which was sampled and seeded for the experiment. Similarly, 1.24 × 10^5^ barcoded HepG2 cells were expanded (×102) to form the POT, which was sampled and seeded for the experiment.

### Treatment-recovery of barcoded cell lines

Barcoded HuH6 cells were seeded into 15 T175 flasks: three were treated with NaCl, the remaining 12 were treated with cisplatin (20μM; C3374, Cambridge Bioscience, Cambridge, UK). After 48 hours, three flasks were sampled, and the remaining cultures were washed three times (1× PBS, 2× DMEM) to remove the drug. Preliminary experiments were used to inform both on a dose that would cause a bottleneck but that could also be recovered from. For the HuH6, preliminary experiments suggested a full recovery took place after three months. Therefore, recovery samples were taken in triplicate at 7-, 24- and 91-days post-treatment. Samples were cryopreserved for 10x Genomics (Pleasanton, CA, USA) scRNA-seq or frozen down as cell pellets for DNA barcode sequencing.

Similarly, barcoded HepG2 cells were seeded into 12 T75 flasks: three were treated with NaCl, nine were treated with cisplatin (8μM; Cambridge Bioscience). After 48 hours, three flasks were sampled, and remaining cultures were washed three times. As with the HuH6 cell line, preliminary experiments were used to estimate doses suitable for long-term barcode experiments and fully recovery, which for this cell line was around 1 month – recovery samples were taken in triplicate at 10- and 35-days post-treatment. Samples were cryopreserved for 10x Genomics scRNA-seq or frozen down as cell pellets for DNA barcode sequencing.

### DNA barcode sequencing

Barcode amplification by PCR was performed using Titanium Taq (639208, Takara Bio, Kusatsu, Japan), 1500ng of the extracted DNA, the forward XP primer (5’-ACCGAACGCAACGCACGCA-3’; 10μM) and the reverse XP primer (5’-ACGACCACGACCGACCCGAACCACGA-3’; 10μM). NGS libraries were prepared from the PCR product using the NEBnext® Ultra™ II DNA Library Prep Kit for Illumina® (E7645L, New England Biolabs, Ipswich, MA, USA) and NEBNext® Multiplex Oligos for Illumina® (E6442S, New England Biolabs). NGS was performed at the Genomics Facility using the Illumina (San Diego, CA, USA) NovaSeq 6000. 150 pair-end reads and 5% PhiX was used and approximately 5000 reads per time-point was targeted.

### DNA barcode analysis

Paired FASTQ files were mapped to a custom index reference, generated from all possible sequences of Cellecta barcodes, using Burrows-Wheeler Aligner v0.7.17 (Li & Durbin, 2009). The BAM files were generated using Sequence Alignment/Map v1.18 (Li *et al*, 2009). Gene counts were quantified using the featureCounts function from Rsubread v2.20.0 (Liao *et al*, 2019).

### scRNA-seq of barcoded cell lines

6000-7500 cells per sample were multiplexed using feature barcoding technology for cell multiplexing from 10x Genomics and taken into the 10x reactions using the Chromium Next GEM Single Cell 3’ Reagent Kits v3.1 (1000269, 10x Genomics) and prepared for Illumina sequencing on a NovaSeq with S1 flow cell.

### scRNA-seq analysis of barcoded cell lines

10x Genomics Cell Ranger v6.1.2 (Zheng *et al*, 2017) was used to process raw sequencing FASTQ files. EmptyDrops from DropletUtils v1.20.0 was used to remove empty droplets with a false discovery rate of 0.1%.

Cell-level filtering was carried out with the following thresholds: mitochondrial gene content <10%, ribosomal gene content >5%, and detected genes >1000 (Appendix Figure S3, S4; Appendix Table S1). Filtering at the gene-level was implemented to remove genes without a HGNC ID and those with fewer than five reads.

The NormalizeData function from the Seurat v5.0.0 (Butler *et al*, 2018) package was applied, FindVariableFeatures was used to identify highly variable genes (HVGs), which were filtered to remove mitochondrial, long non-coding and ribosomal genes, and the data was scaled using ScaleData. A principal component analysis (PCA) was performed on the normalised-scaled data. The batch correction algorithm Harmony v1.2.3 (Korsunsky *et al*, 2019) was applied, and the output used for uniform manifold approximation and projection (UMAP)-based dimensional reduction.

The data was clustered using FindClusters at resolutions from 0.2 to 0.8 (in increments of 0.2) for the whole dataset and the data split by experimental timepoint (Condition). FindAllMarkers was used to obtain cluster markers and pairwise Jaccard similarity was carried out to compare all the clusters in the whole and split datasets. The silhouette function from cluster v2.16 (Struyf *et al*, 1996) was used to compute silhouette information for *k* = 2 to *k* = 15 to inform the value of *k* to assign when running the function pam on the entire dataset. Markers for PAM clusters were found using Seurat’s FindAllMarkers. Annotation of clusters was carried out referencing HB signatures (Cairo *et al*, 2008; Carrillo-Reixach *et al*, 2020b; Roehrig *et al*, 2024b), liver development signatures (Wesley *et al*, 2022), and liver development markers (Zong & Stanger, 2012; Lemaigre, 2009; Si-Tayeb *et al*, 2010) taken from the literature (Appendix Table S2). HuH6 expression clusters were found to fall into three larger expression states, Hep-/Pro-/Stem-like and a differential gene expression analysis was carried out between these three states to find markers of each (Supplementary Data 1). These gene sets were used in Figure EV1F to compare HuH6 and HepG2 expression clusters. Hep-like genes were also used in Figure EV3B to identify expression state genes within the (HuH6-derived) recovery-up gene list.

Pseudotime analysis was carried out using the R package Slingshot v2.14.0 (Street *et al*, 2018) on the PCA results. The root was set to cluster 5 (Pro-like) for both the HuH6 and HepG2 datasets based on preliminary understanding of recovery from cisplatin. Gene dynamics analysis by fitting a linear spline model to each Slingshot lineage was carried out using TSCAN v1.44.0 (Ji & Ji, 2016). Gene set enrichment analysis (GSEA) for Gene Ontology Biological Processes and MSigDB C2 and Hallmark gene sets was performed using clusterProfiler v4.14.6 (Yu *et al*, 2012) and msigdbr v7.5.1 (Dolgalev, 2018).

Signature scoring of single cells was carried out using Seurat’s AddModuleScore for Hallmark EMT and G2M gene lists. Spearman rank correlation coefficients were calculated, the *p*-values for which are also noted but all reached R’s lower threshold of reporting, likely driven by large numbers of cells.

Enrichment analysis of transcription factor binding motifs in gene lists derived from the pseudotime analysis was carried out using RcisTarget v4.14.6 (Sara Aibar, 2018). Gene-motif rankings databases were downloaded from the cisTarget resources website (https://resources.aertslab.org/cistarget/). HGNC motif annotations were loaded, and the cisTarget function was used for motif enrichment analysis using the 500bp upstream TSS database.

### Single-nuclei multiome sequencing (snRNA-seq and snATAC-seq) of barcoded cell lines

We performed single-nuclei multiome sequencing (snRNA-seq and snATAC-seq) on two untreated and two recovered (Recovery t_3_) HuH6 replicates. Samples were prepared for sequencing following the 10x Genomics Multiome (snRNA-seq and snATAC-seq) protocol. Samples were processed targeting 2400-6000 nuclei per sample following the 10x Genomics Next GEM Single Cell Multiome ATAC + Gene Expression protocol and prepared for Illumina sequencing on a NovaSeq with S1 flow cell.

### Single-nuclei mutliomics analysis of barcoded HuH6

We used Cell Ranger ARC v2.0.0 (Satpathy *et al*, 2019) to align snRNA-seq and snATAC-seq reads to the human genome (GrCh38/hg38) and generate the matrices of UMI counts per gene and ATAC-seq fragments.

As described in *scRNA-seq analysis of barcoded cell lines*, cell-level filtering was carried out with the following thresholds: mitochondrial gene content <30% and detected genes >800. Filtering at the gene-level was implemented to remove genes without a HGNC ID and those with fewer than five reads. The RNA data was normalised, scaled, and reduced by PCA. Harmony batch correction was applied, and the output used for UMAP-embedding.

Clustering followed the approach outlined in *scRNA-seq analysis of barcoded cell lines*: FindClusters across resolutions from 0.2 to 0.8 were tested for the whole dataset and the data split by experimental timepoint (Untreated, Recovered); FindAllMarkers was used to obtain cluster markers and pairwise Jaccard similarity was carried out to compare all the clusters in the whole and split datasets; silhouette (*k* = 2 to *k* = 15) was used to inform the value of *k* to assign when running pam. As in *scRNA-seq analysis of barcoded cell lines*, clusters were annotated by their expression of HB signatures (Cairo *et al*, 2008; Carrillo-Reixach *et al*, 2020b; Roehrig *et al*, 2024b), liver development signatures (Wesley *et al*, 2022), and liver differentiation markers (Lemaigre, 2009; Si-Tayeb *et al*, 2010; Zong & Stanger, 2012), and their similarity to previously defined scRNA-seq clusters.

For the ATAC data, quality control (QC) filtering was applied ad hoc for each sample on the following metrics: fragment counts, UMI counts, nucleosome signal, and transcription start site (TSS) enrichment. Peak calling was carried out using the function CallPeaks, which relies on MACS v3.0.3 (Zhang *et al*, 2008) peak calling, from Signac v1.14.0 (Stuart *et al*, 2021) on each sample separately. Blacklisted peaks and peaks on sex chromosomes were removed. The peak sets were then combined with the reduce function from GenomicRanges v1.58.0 (Lawrence *et al*, 2013), which merges intersecting peaks. The data was merged into a single Signac object with the combined peak set.

The resulting peak-cell count matrix was normalised by the term frequency-inverse document frequency normalisation procedure (Cusanovich *et al*, 2015). FindTopFeatures was applied to compute the feature metadata and RunSVD employed for singular value decomposition to return a reduced dimension representation of the object. UMAP dimensionality reduction for visualisation was applied. Seurat functions FindNeighbours and FindClusters were used to identify clusters between resolutions from 0.2 to 0.8. Differential accessibility analysis was carried out between recovered and untreated conditions on target genes.

### Patient samples and datasets

Four patient samples were obtained from VIVO biobank, Newcastle, UK. These samples were matched pairs from two patients: patient 1 mid-treatment and post-treatment relapse samples; patient 2 post-treatment and post-treatment relapse samples. Informed consent was obtained from all patients, or from their parents or legal guardians where appropriate, for the collection, storage and research use of these samples. Samples were provided under VIVO Biobank’s ethical approval. All experiments conformed to the principles set out in the WMA Declaration of Helsinki and the Department of Health and Human Services Belmont Report. Expression data was obtained from snRNA-seq of frozen tissue samples (see below).

Patient data analysed from Chen *et al*. (2025) were longitudinal single-cell profiles from pre- and post-chemotherapy samples, available for five patients, which were included in their larger cohort study. This data is available via NCBI’s Gene Expression Omnibus under the accession code GSE326150 (Chen *et al*, 2026).

The patient dataset from Roehrig *et al*. (2024) was composed of six HB samples and five NT liver, collected from five patient, originally part of a larger cohort (Hirsch *et al*, 2021). Data is accessible via the European Genome Archive under the accession code EGAS00001006932 (Roehrig *et al*, 2024a).

The discovery set described by Carrillo-Reixach *et al*. (2020) included 65 samples from 33 patients (31 primary tumours, 2 recurrences and 32 paired adjacent NT liver) were taken. Expression data was obtained through RNA-seq of 64 frozen tissue samples form 32 patients (32 HB and 32 paired NT). This dataset can be accessed through NCBI’s Gene Expression Omnibus under the accession code GSE132219 (Carrillo-Reixach *et al*, 2020a).

### snRNA-seq of patient samples

Longitudinal pairs of clinical samples were processed for snRNA-seq. Briefly, nuclei were isolated by mincing samples in a Tween-based lysis buffer containing RNase inhibitors, washed, counted on a LUNA fluorescence cell counter (Logos Biosystems, Gyeonggi-do, South Korea) and prepared for the 10x Chromium using the Chromium Next GEM Single Cell 3ʹ Kit v4 (10x Genomics), targeting 20,000 nuclei per sample. cDNA and libraries were prepared for sequencing with the same Chromium kit and sequenced (PE150) on a NovaSeq X (Illumina).

### snRNA-seq analysis of patient samples

As outlined in *scRNA-seq analysis of barcoded cell lines*, raw FASTQ files were processed with Cell Ranger v6.1.2 (Zheng *et al*, 2017), and empty droplets were excluded using EmptyDrops from DropletUtils v1.20.0, with a false discovery rate of 0.1%.

Cell-level filtering was carried out with the following thresholds: mitochondrial gene content <15% and detected genes >1000 (Appendix Figure S5; Appendix Table S3). Filtering at the gene-level was implemented to remove genes without a HGNC ID and those with fewer than five reads.

Data was normalised and scaled following the same approaches described in *scRNA-seq analysis of barcoded cell lines*. PCA and Harmony-based batch correction were applied and used for UMAP embedding.

FindClusters across resolutions of 0.2 to 0.8 were tested for the pooled dataset and the data split by patient (Patient_ID). FindAllMarkers and pairwise Jaccard similarity were used to compare clusters. Silhouette scores were evaluated across *k* = 2 to *k* = 30. PAM cluster markers were obtained via FindAllMarkers. Clusters were annotated using HB signatures (Cairo *et al*, 2008; Carrillo-Reixach *et al*, 2020b; Roehrig *et al*, 2024b), liver development signatures (Wesley *et al*, 2022), and liver differentiation markers (Lemaigre, 2009; Si-Tayeb *et al*, 2010; Zong & Stanger, 2012). Cluster annotation was also completed with SingleR v2.8.0 analysis using the Human Primary Cell Atlas to annotate cell types. InferCNV v1.21.0 was used to aid identification of malignant cells, with default parameters and immune cells as the reference.

### Patient data statistical analysis

The statistical analysis on data from Carrillo-Reixach *et al*. (2020) was performed with SPSS (v26) and GraphPad Prism (v10.1.0) software. Appropriate two-side statistical tests were applied when necessary for qualitative variables: chi-square and Fisher’s exact tests are indicated in each analysis. We applied the Wilcoxon matched-pairs signed-rank test for non-parametric data to compare two different groups. The Shapiro-Wilk test was performed for the study of the normality of the data. For EFS analysis, Kaplan-Meier’s method and log-rank test were performed to compare differences between curves.

### BIRC5 siRNA knockdown

HuH6, HepG2, and HB303 cells were plated in 6-well plates (1 ×10^5^ cells/well for HuH6 and 2 × 10^5^ cells/well for HB303 and HepG2). Three wells were transfected with a non-targeting control (NTC; D-001810-10-05, Horizon Discovery, Cambridge, UK) and three wells were transfected with BIRC5 siRNA (L-003459-00-0005, Horizon Discovery). Transfection reagent was removed 6 hours after addition and replaced with complete media. 72 hours post-transfection cells were harvested, and two wells were viably frozen for scRNA-seq, while one well was flash frozen as a cell pellet for western blot analysis.

Protein expression was assessed by western blotting using cell lysates (30µg). Cell lysates were resolved on a Bolt™ Bis-Tris gel and transferred on a PVDF membrane. Primary antibodies, BIRC5 (2808T, Cell Signalling, Danvers, MA, USA) and GAPDH (2118S, Cell Signalling), were incubated overnight. After incubation with the secondary antibody for 1 hour, proteins were detected by enhanced chemilumiescence using SuperSignal™ (34580, ThermoFisher, Waltham, MA, USA).

Duplicates per condition were sequenced by scRNA-seq. 10,000 cells per sample were multiplexed and put into the 10x reactions using the Chromium Next GEM Single Cell 3’ Reagent Kits v3.1 (1000269, 10x Genomics) and prepared for Illumina sequencing on a NovaSeq with S1 flow cell.

Analysis of *BIRC5* siRNA knockdown scRNA-seq data was carried out as outlined in *scRNA-seq analysis of barcoded cell lines*, i.e. Cell Ranger and Seurat workflow – filtering, normalisation, scaling, clustering, cluster annotation. Filtering thresholds were as follows: mitochondrial gene content <20%, ribosomal gene content >5%, and detected genes >1000 (Appendix Figure S6; Appendix Table S7); genes with >5 reads retained.

### YM155 drug response assays

HuH6, HepG2, and HB303 were plated in 96-well optical-bottom microplates. HB7 T-org and HB22 T-org cells were plated in 96-well plates previously coated with BME matrigel. At cell seeding point, 10^4^ cells were seeded per well and the experiments started when confluence reached 40-50%. Three wells were treated per condition with the control (media), vehicle (DMSO), or eight treatment points: for cell lines – 1nM, 2nM, 5nM, 10nM, 20nM, 50nM, 100nM, 200nM diluted in their respective culture media; for PDO cultures – logarithmically distributed from 0.001 to 2µM diluted in BEC media. Response curves were obtained after two days of treatment for cell lines and after four days (with one medium refresh day two) of treatment. The readout of the experiments was obtained using CellTiter-Glo (G9683, Promega, Madison, WI, USA), which measures the presence of ATP by luminescence. The curves were obtained through a nonlinear regression model in R.

### Cisplatin-YM155 sequential treatment on PDO cultures

HuH6, HepG2, and HB303 were plated in 96-well optical-bottom microplates. HB7 T-org and HB22 T-org cells were plated in 96-well plates previously coated with BME matrigel. At cell seeding point, 10^4^ cells were seeded per well and the experiments started when confluence reached 40-50%. All lines were treated under the same protocol (six wells/condition). At the start of the experiment, half of the plate was treated with cisplatin (IC50 for cell lines; 25µM for PDO cultures). After 72 hours, when the cells were visually affected by cisplatin, all cells were washed twice with cooled PBS and treated with the following conditions: maintenance of media, maintenance of cisplatin, 0.01µM YM155, 0.2µM YM155, or (PDO cultures only) 0.6µM YM155. After a further 72 hours of sequential treatment, CellTitre-Glo readout was performed.

## Data availability

The raw sequencing data used and generated during this study has been deposited in the National Centre for Biotechnology Information – Sequence Read Archive (NCBI-SRA) under the accession number PRJNA1247955. Processed data has been deposited on Zenodo (doi: 10.5281/zenodo.18889051). The scripts used to analyse the data for this study are available on GitHub (https://github.com/PCEBrunaLab/Chen-et-al-2025).

## Author contributions

Conceptualisation – AB; data curation – ASS, EC; formal analysis – AMCM, EC, MMM, LZ, SH; funding acquisition – AB, CA, PSB; investigation – CE, EC, MMM, LZ; methodology – CE, EC, MMM, LZ; project administration – AB; resources – AB, CA, EI, PSB, SC; software – ASS, EC; supervision – AB; validation – CE, EC; visualisation – AAA, AMCM, CE, EC, MMM, LZ; writing – original draft – AB, CE, EC; writing – review and editing – AAA, AB, AMCM, CA, CE, EC, MMM, LZ, PSB, SC, SH.

## Acknowledgements

We thank Yura Grabovska (Institute of Cancer Research, London, UK) for insightful bioinformatic suggestions and discussions; Theo Hirsch (Centre de Recherche des Cordeliers, Paris, France) for assistance with data access; Roland Kappler (Ludwig-Maximilians-Universität, Munich, Germany) for providing the HuH6 cell line; and Peijie Zhou (University of California, Irvine, USA) for guidance on implementing MuTrans. We would also like to acknowledge the contributions of the ICR Core Research Facilities – Flow Cytometry (https://doi.org/10.5281/zenodo.14845954), Genomics (https://zenodo.org/records/14845894) and Scientific Computing.

This work is supported by: Grace Kelly Childhood Cancer Trust – AB2025GKCCT (AB); Scientific Foundation of the Spanish Association Against Cancer – IGTP-IDIBAPS-AECC 2022-042/PRYCO223102ARME (CA, PSB); Fight Kids Cancer Funding Programme supported by Imagine for Margo, Fondation KickCancer, Foundation Kriibskrank Kanner, Federazione Italiana Associazioni Genitori e Guariti Oncoematologia Pediatrica, Cris Cancer Foundation – BT4ChildLC (CA); Agency for Management of University and Research Grants AGAUR – 2021-SGR-01186 (CA); Worldwide Cancer Research – 23-0356 (PSB). Patient samples and data used in this study were provided by VIVO Biobank, supported by Cancer Research UK & Blood Cancer UK – CRCPSC-Dec21\100003.

---

**Figure EV1:**
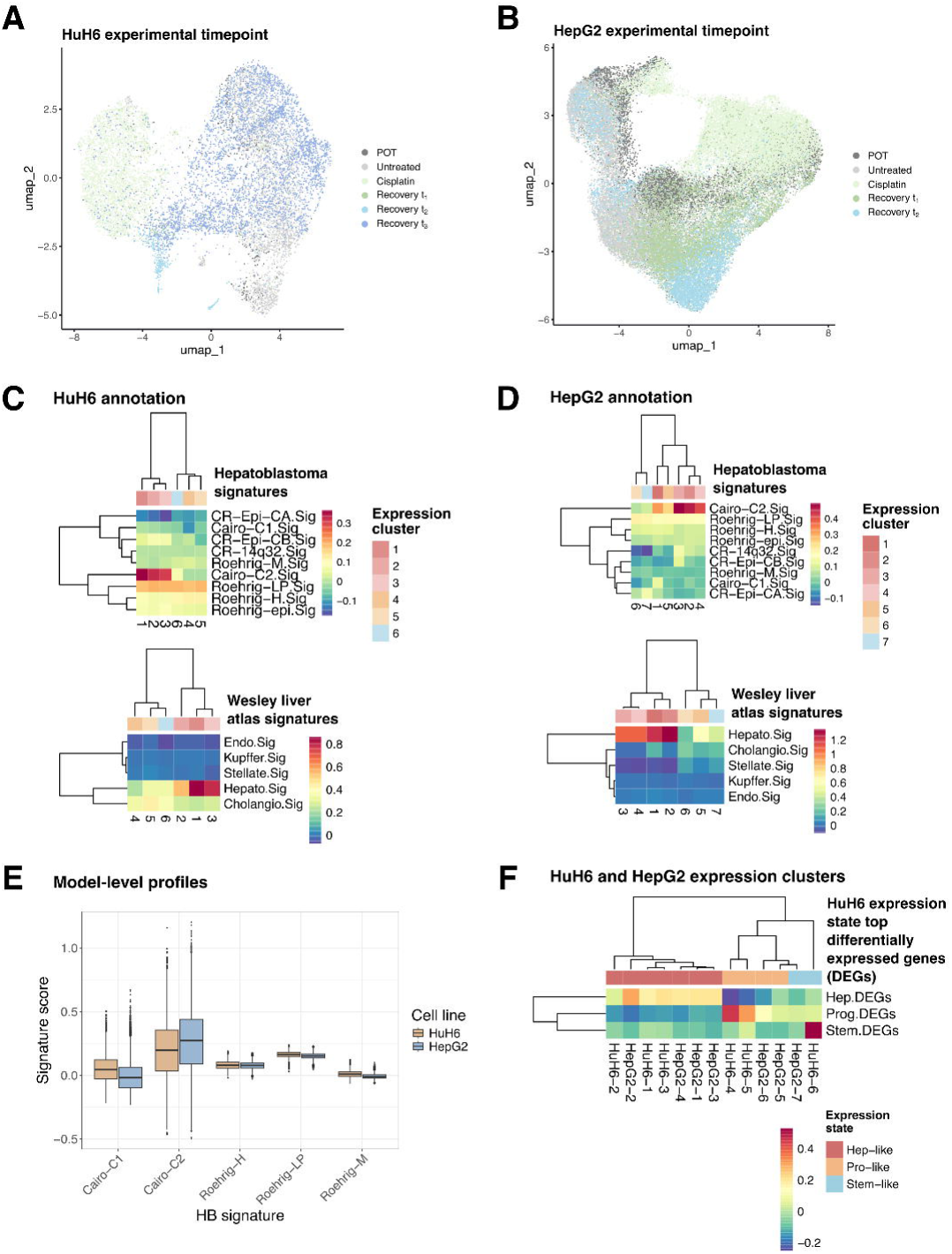
**(A)** UMAP of 10,791 HuH6 cells that passed QC, coloured by experimental timepoint. **(B)** UMAP of 43,017 HepG2 cells that passed QC, coloured by experimental timepoint. **(C)** Signature scoring of HuH6 expression clusters for HB signatures (Cairo et al, 2008; Carrillo-Reixach et al, 2020b; Roehrig et al, 2024b) and liver development signatures (Wesley et al, 2022). **(D)** Signature scoring of HepG2 expression clusters for HB signatures (Cairo et al, 2008; Carrillo-Reixach et al, 2020b; Roehrig et al, 2024b) and liver development signatures (Wesley et al, 2022). **(E)** Signature scoring of HuH6 and HepG2 cells (at the model-level) for Cairo C1/C2 (Cairo et al, 2008) and Roehrig H/LP/M (Roehrig et al, 2024b) signatures. **(F)** Expression of HuH6-derived differentially expressed genes between the Hep-/Pro-/Stem-like expression states in HuH6 and HepG2 scRNA-seq data.

**Figure EV2:**
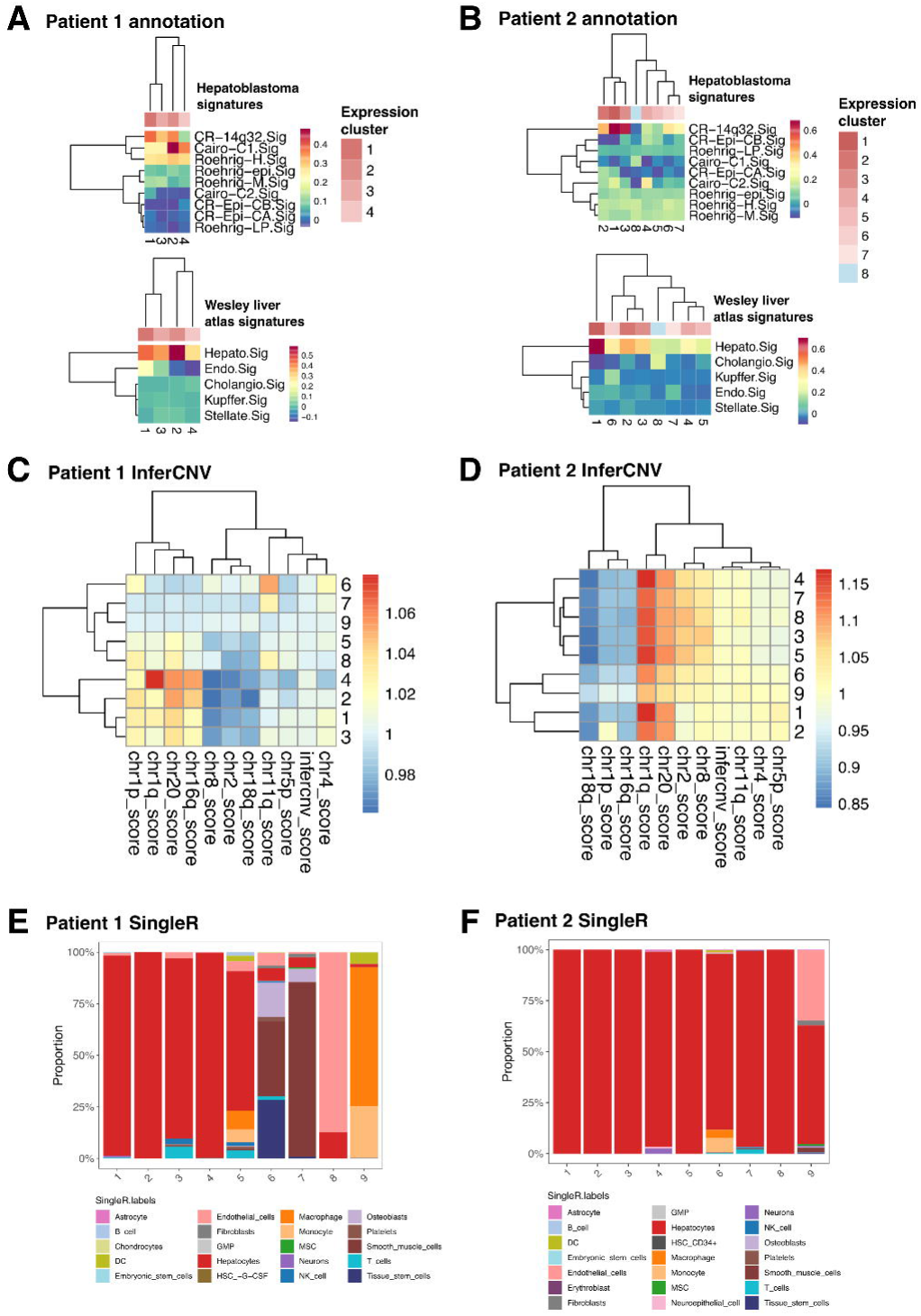
**(A)** Signature scoring of Patient 1 (P1) tumour expression clusters (1-4) for HB signatures (Cairo et al, 2008; Carrillo-Reixach et al, 2020b; Roehrig et al, 2024b) and liver development signatures (Wesley et al, 2022). **(B)** Signature scoring of Patient 2 (P2) tumour expression clusters (1-8) for HB signatures (Cairo et al, 2008; Carrillo-Reixach et al, 2020b; Roehrig et al, 2024b) and liver development signatures (Wesley et al, 2022). **(C)** InferCNV scoring of P1 all expression clusters (1-9) for commonly altered chromosome regions in HB. **(D)** InferCNV scoring of P2 all expression clusters (1-9) for commonly altered chromosome regions in HB. **(E)** Proportion of cells annotated by SingleR cell types for P1 expression clusters. **(F)** Proportion of cells annotated by SingleR cell types for P2 expression clusters.

**Figure EV3:**
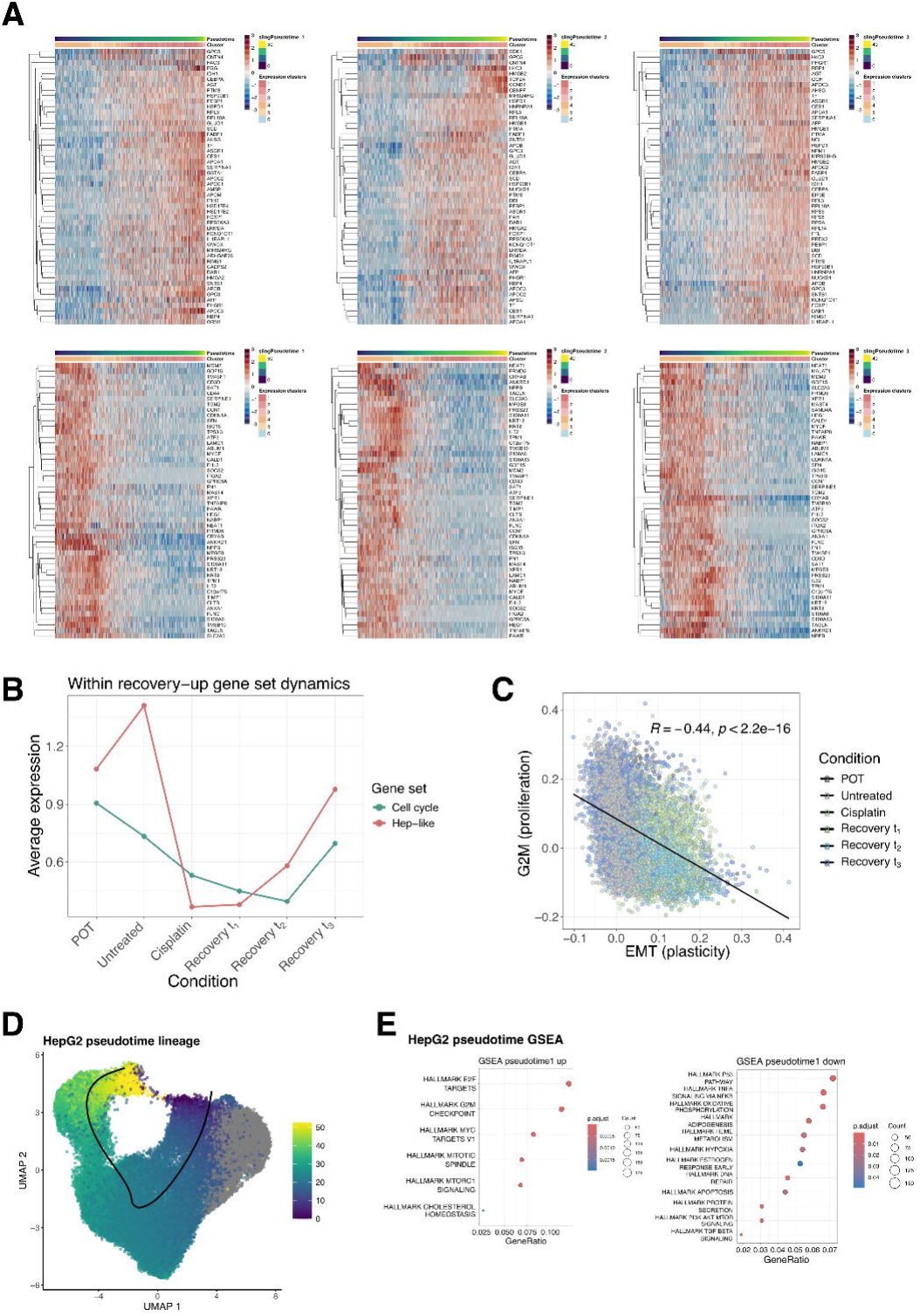
**(A)** Heatmaps showing the HuH6 scRNA expression of the top 50 genes, as ranked by log_2_ fold change, whose expression increased or decreased with pseudotime progression along recovery. Each panel from left-to-right represents a Slingshot pseudotime progression: recovery lineage 1, recovery lineage 2, and recovery lineage 3. Top-to-bottom the panels are arranged as (top) heatmaps showing genes that increase in expression as pseudotime progresses and (bottom) heatmaps showing genes that decrease in expression as pseudotime progresses. Cells (columns of each heatmap) were ordered according to pseudotime value. **(B)** The average expression across experimental timepoint of Hep-like genes (derived from HuH6 differential gene expression analysis between Hep-/Pro-/Stem-like states; see Methods and Supplementary Data 1) and cell cycle genes in recovery-up gene list. **(C)** Correlation between cells signature scored by Hallmark EMT and Hallmark G2M gene sets. Coloured by experimental timepoint. Spearman’s rank correlation coefficient, *r_s_* = −0.44 (*p* < 2.2 x 10^-16^; *p*-value is R’s lower reporting threshold, common for single-cell data as large numbers of cells results in high power tests). **(D)** scRNA-seq UMAP of 43,017 HepG2 cells that passed QC, coloured by computed shared pseudotime value. Black line represents recovery lineage. **(E)** GSEA plots for MSigDB Hallmark terms that are associated with the genes in the HepG2 pseudotime analysis whose expression, either increasing (up) or decreasing (down), correlates with pseudotime progression (pseudotime-derived genes).

**Figure EV4:**
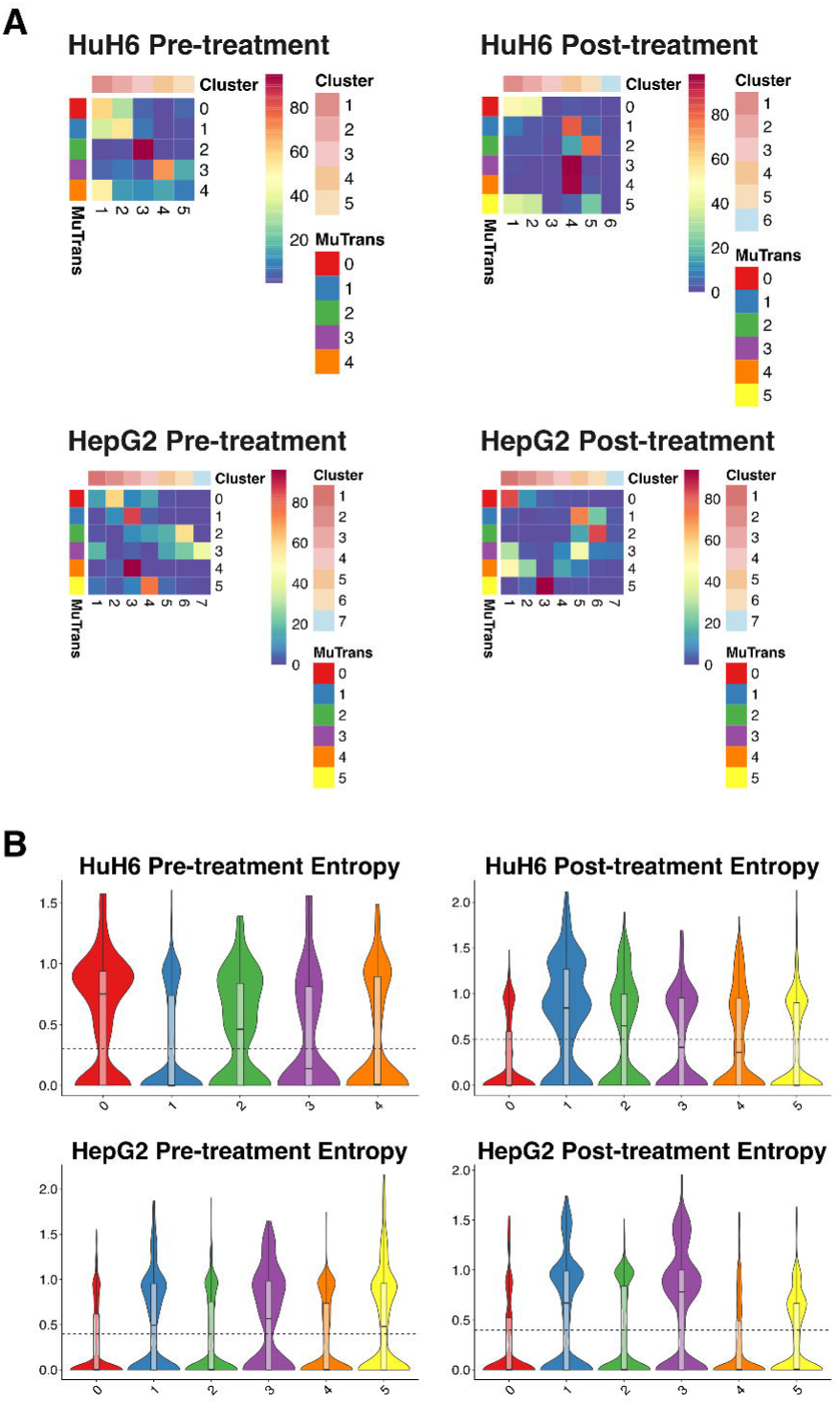
**(A)** Number of cells that occupy a given MuTrans attractor wells and expression cluster for each of the pre- and post-treatment datasets for the HuH6 and HepG2 scRNA-seq datasets. **(B)** Violin plots of the entropy of HuH6 and HepG2 cells predicted by MuTrans and the estimated cutoff threshold between high and low entropy cells (dashed line).

**Figure EV5:**
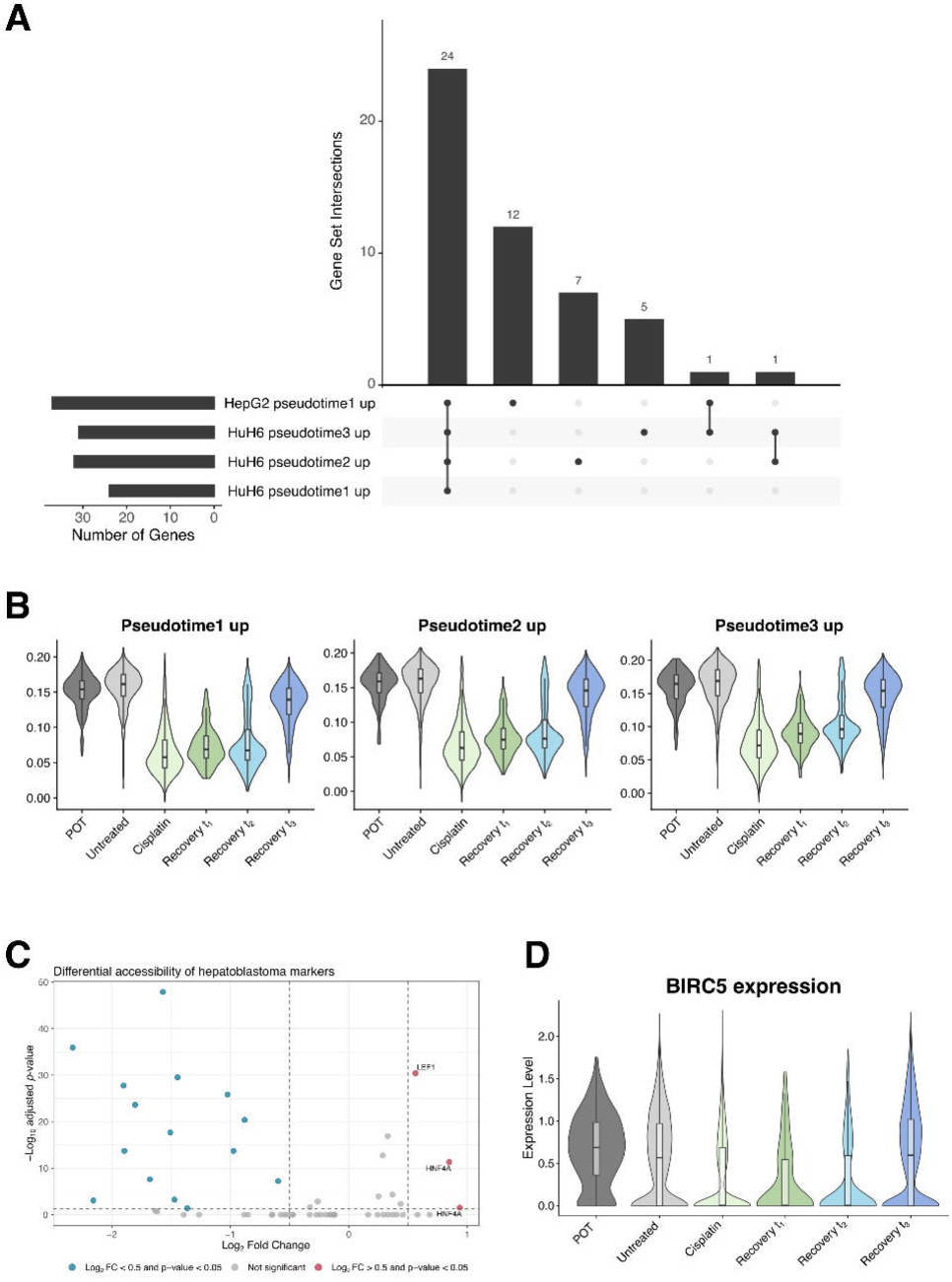
**(A)** Overlap between transcription factor motifs enriched in genes that increase in expression as pseudotime increases for each of the HuH6 recovery lineages (1-3) and the HepG2 recovery lineage. **(B)** Expression of recovery-up markers at each experimental timepoint in the HuH6 scRNA-seq data. **(C)** Differential accessibility analysis between recovered and untreated samples of known HB markers (*AFP*, *CTNNB1*, *HNF4A*, *LEF1*, *TCF4*), with thresholds of log_2_ fold change > 0.5 and adjusted *p*-value < 0.05. Each point represents a region, and multiple regions may be associated with the same gene, hence multiple points may be labelled with the same gene. **(D)** Expression of *BIRC5* at each experimental timepoint in the HuH6 scRNA-seq data.

**Figure EV6:**
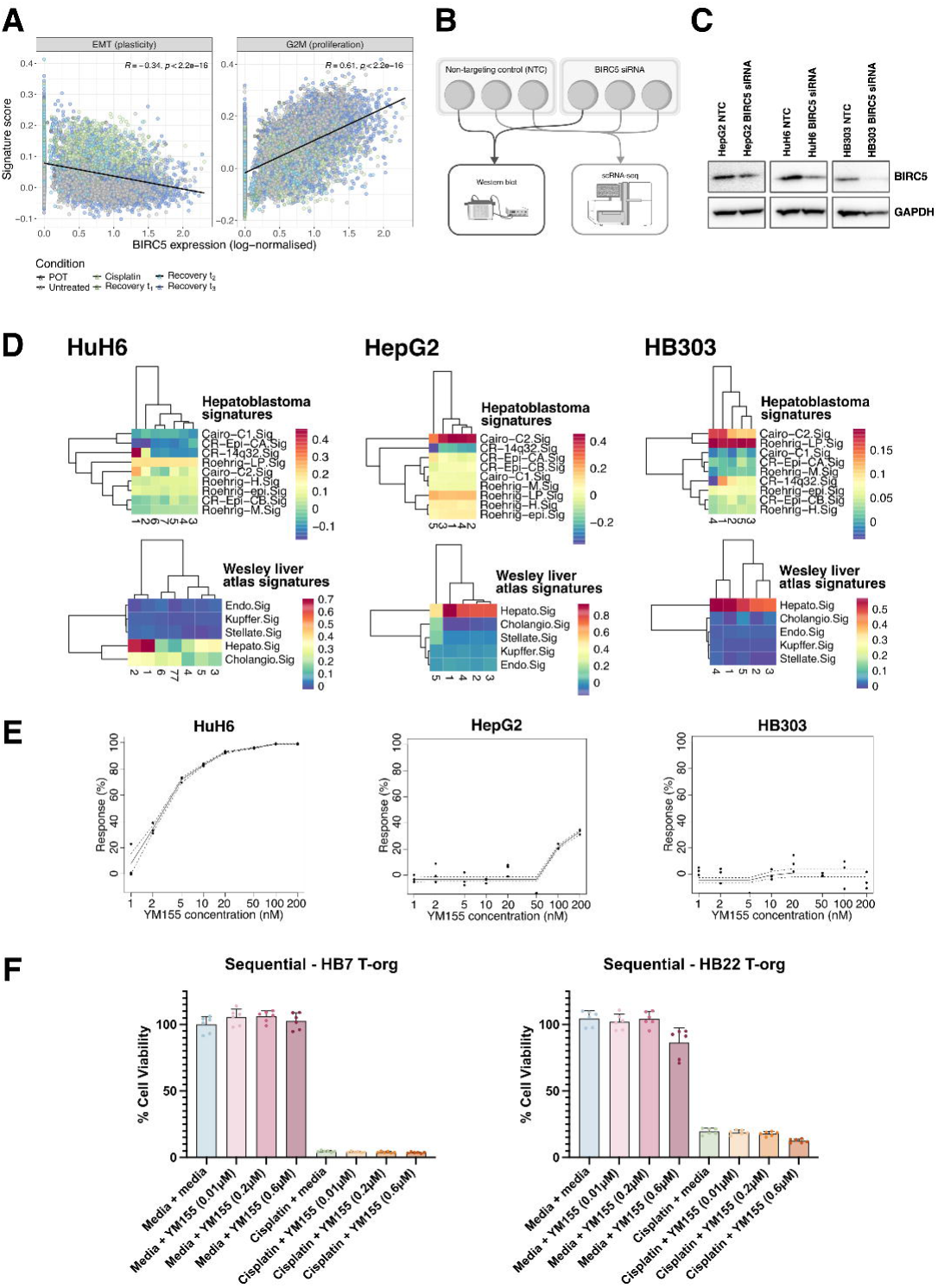
**(A)** Correlation between the *BIRC5* expression of cells and their signature score for either Hallmark EMT (left) and Hallmark G2M (right) gene sets. Coloured by experimental timepoint. Spearman’s rank correlation coefficient for *BIRC5*-EMT, *r_s_* = −0.34 and *BIRC5*-G2M, *r_s_* = 0.61 (both *p* < 2.2 x 10^-16^; *p*-value is R’s lower reporting threshold, common for single-cell data as large numbers of cells results in high power tests). **(B)** *BIRC5* siRNA knockdown experimental design. **(C)** Western blot analysis of BIRC5 protein in HuH6, HepG2, and HB303 cells transfected with a non-targeting control (NTC) and *BIRC5* siRNA. **(D)** Signature scoring for transfected cell line samples – (left-to-right) HuH6, HepG2, HB303 – pseudobulked by expression cluster, for HB signatures (Cairo et al, 2008; Carrillo-Reixach et al, 2020b; Roehrig et al, 2024b) and liver development signatures (Wesley et al, 2022). **(E)** Drug-response curves in (left-to-right) HuH6, HepG2, HB303 for YM155 (concentration ranges from 1 to 200nM). **(F)** Sequential cisplatin treatment with media or YM155 at three concentrations (0.01µM, 0.2µM, 0.6µM), and media + YM155 controls, unnormalized cell viability values.

