## Supplementary material for "Integrated Lineage Tracing in Hepatoblastoma Finds a Plasticity-Proliferation Axis Drives Adaptation": Table 1; Table 2

**Table 1 – Patient metadata**

| **Sample Name** | **Gender** | **Time to T2** | **Chemo** | **Location** | **Histology** | **Tumour Fraction** | **Ploidy** |
| --- | --- | --- | --- | --- | --- | --- | --- |
| Patient1 T1 | Male | 18 months | Mid | Liver | Epithelial – foetal | 0.6552 | 1.9988 |
| Patient1 T2 | Male |  | Post-relapse | Liver | Epithelial – foetal | 0.3388 | 2.0327 |
| Patient 2 T1 | Female | 5 months | Post | Liver | Mixed – mostly epithelial | 0.8088 | 1.9585 |
| Patient 2 T2 | Female |  | Post-relapse | Lymph node | Mixed | 0.7939 | 1.9821 |

**Table 1**: Patient sample metadata retrieved from pathology reports and low-pass whole genome sequencing.

**Table 2 – *BIRC5* expression in clinical samples**

|  | | ***BIRC5* gene expression** | |
| --- | --- | --- | --- |
|  |  | ***BIRC5* < median** | ***BIRC5* ≥ median** |
| Age ≥ 8 years | No | 15 | 13 |
|  | Yes | 1 | 3 |
|  | *p*-value | 0.3 | |
| Metastasis | No | 12 | 11 |
|  | Yes | 4 | 5 |
|  | *p*-value | 0.5 | |
| Vascular Invasion | No | 11 | 8 |
|  | Yes | 5 | 8 |
|  | *p*-value | 0.280* | |
| Multifocality | No | 11 | 9 |
|  | Yes | 5 | 7 |
|  | *p*-value | 0.465* | |
| PRETEXT-4 | No | 12 | 14 |
|  | Yes | 3 | 2 |
|  | *p*-value | 0.468 | |
| CHIC-HS | VLR/LR/IR | 11 | 9 |
|  | HR | 5 | 7 |
|  | *p*-value | 0.465* | |
| Histology | Epithelial | 6 | 10 |
|  | Mixed | 10 | 6 |
|  | *p*-value | 0.157* | |
| Main Epithelial Component | Foetal | 11 | 8 |
|  | Non-foetal | 4 | 8 |
|  | *p*-value | 0.168 | |
| Epi-CB Subtype | No | 12 | 6 |
|  | Yes | 4 | 10 |
|  | *p*-value | 0.037 | |
| 14q32 Amplification | Moderate | 10 | 2 |
|  | Strong | 6 | 14 |
|  | *p*-value | 0.005 | |
| C2-Pure Subtype | No | 15 | 6 |
|  | Yes | 1 | 10 |
|  | *p*-value | 0.001 | |
| Molecular Risk Stratification | MRS-1/2 | 13 | 6 |
|  | MRS-3 | 3 | 10 |
|  | *p*-value | 0.015 | |

**Table 2**: *BIRC5* gene expression with clinical – age, presence of metastasis, vascular invasion, multifocality, PRE-Treatment EXTension of disease (PRETEXT), and Children’s Hepatic Tumours International Collaboration-Hepatoblastoma Stratification (CHIC-HS); pathological – histology and main epithelial component; and molecular variables – 14q32 amplification-gene signature expression, epigenetic Epi-CA/CB subtypes, C1/C2-Pure/C2B transcriptomic classification, and Molecular Risk Stratification (MRS). The main epithelial component, as described by Cairo *et al.* (2008) is classified as either “foetal” (more than 50% of tumour cross-sectional areas are sheets of uniform, cuboidal cells showing low mitotic activity) or “non-foetal” (other predominant components, different from foetal, that may include crowded foetal macrotrabecular and embryonal histological patterns). Chi-square* and Fisher’s exact tests were used as appropriate.
