## Appendix Figure SX; Appendix Table SX for "Integrated Lineage Tracing in Hepatoblastoma Finds a Plasticity-Proliferation Axis Drives Adaptation"

Table of Contents

Appendix Figure S1…………………………………………………………………...2

Appendix Figure S2…………………………………………………………………...2

Appendix Figure S3…………………………………………………………………...3

Appendix Figure S4…………………………………………………………………...3

Appendix Figure S5…………………………………………………………………...3

Appendix Figure S6…………………………………………………………………...4

Appendix Table S1…………………………………………………………………......5

Appendix Table S2………………………………………………………………….....5-6

Appendix Table S3…………………………………………………………………......6

Appendix Table S4…………………………………………………………………….6-8

Appendix Table S5……………………………………………………………………8

Appendix Table S6…………………………………………………………………….8-9

Appendix Table S7…………………………………………………………………......9

### Appendix Figures

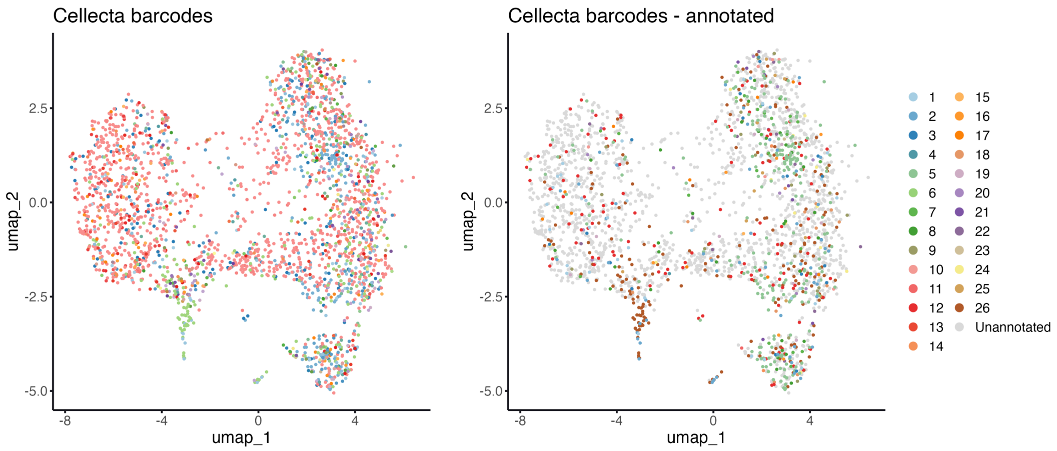


**Appendix Figure S1**: UMAPs of 2,546 HuH6 cells that had both transcriptomic and Cellecta barcode lineage information. (Left) Cells were coloured by their unique barcode identity, of which there were 130 unique barcodes; (right) cells annotated by those that were included in the bubble-pie plots (clones 1-26; Figure 3), i.e. clones for which lineage information was present in the POT sample; clones that did not have ancestors sampled in the POT were grouped as "Unannotated". *N.B.*: Different colours for barcodes 1-26 were used in the plot on the right compared to the plot on the left to differentiate barcodes more clearly.

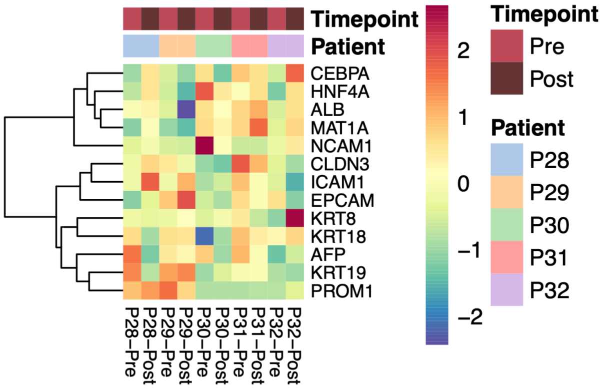


**Appendix Figure S2**: Expression data from Chen *et al.* (2025): 5 patients, pre- and post-chemotherapy. Pseudobulked expression (by patient-sample) of curated panel of liver development markers (see Appendix Table S2). P28, P31, and P32 were all cases that exhibited "partial response", while P29 and P30 showed "stable disease".

### HuH6

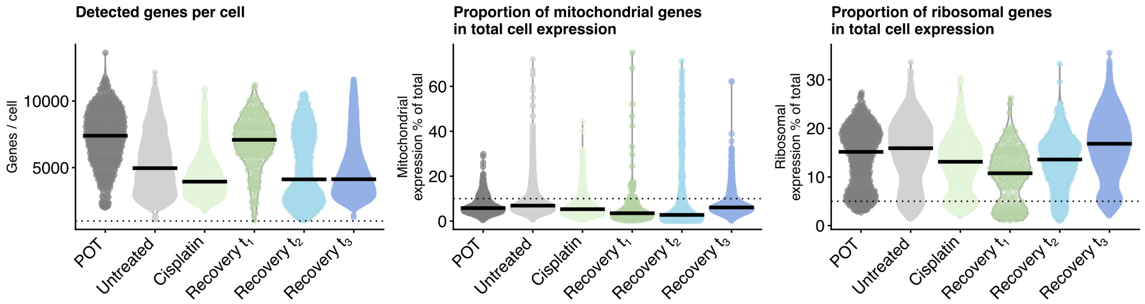


**Appendix Figure S3**: Quality control (QC) features (detected genes, mitochondrial gene content, ribosomal gene content) plotted for the HuH6 barcoding experiment. Filtering thresholds are indicated by dotted lines.

### HepG2

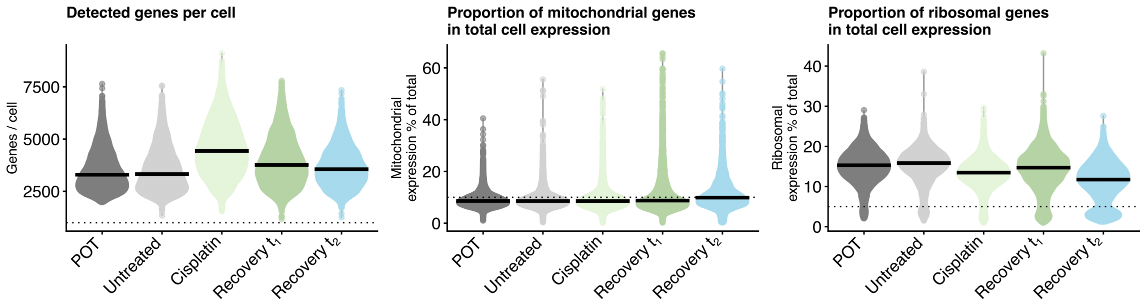


**Appendix Figure S4**: Quality control (QC) features (detected genes, mitochondrial gene content, ribosomal gene content) plotted for the HepG2 barcoding experiment. Filtering thresholds are indicated by dotted lines.

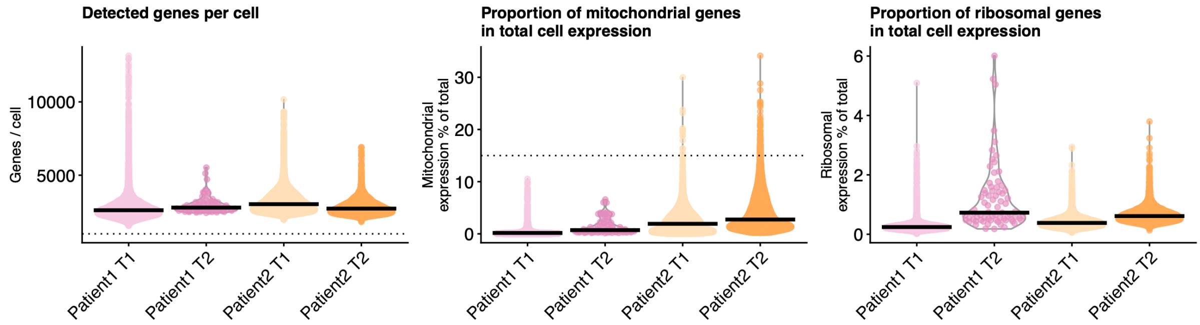


**Appendix Figure S5**: Quality control (QC) features (detected genes, mitochondrial gene content, ribosomal gene content) plotted for the patient case studies. Filtering thresholds are indicated by dotted lines

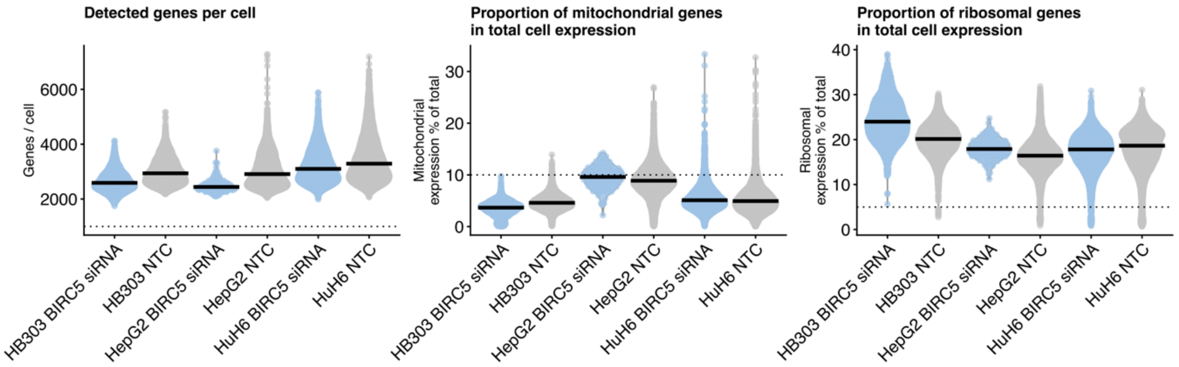


**Appendix Figure S6**: Quality control (QC) features (detected genes, mitochondrial gene content, ribosomal gene content) plotted for the *BIRC5* siRNA knockdown barcoding experiment in three cell lines, HuH6, HepG2, and HB303. Filtering thresholds are indicated by dotted lines. NTC stands for non-targeting control.

**Appendix Tables**

| Condition | HuH6 | HepG2 |
|---|---|---|
| POT | 400 (24.1%) | 8463 (4.6%) |
| Untreated | 2368 (26.6%) | 10623 (6.9%) |
| Cisplatin | 3226 (23.1%) | 12731(6.7%) |
| Recovery $t_1$ | 146 (32.1%) | 7018 (18.9%) |
| Recovery $t_2$ | 372 (29.1%) | 4182 (24.8%) |
| Recovery $t_3$ | 4009 (18.9%) | N/A |

**Appendix Table S1**: Post-filtering cell numbers by condition for HuH6 and HepG2. Values shown are post-filtering cell counts, with the percentage of cells removed during QC filtering shown in brackets.

| Developmental stage | Marker genes | References | Description |
|---|---|---|---|
| Hepatic stem cell | *CLDN3*, *EPCAM*, *NCAM1*, *PROM1* | (Schmelzer *et al*, 2007; Turner *et al*, 2011) | Hepatic stem cells can be distinguished from hepatoblasts by *CLDN3*/*NCAM2* positivity; *EPCAM*/*PROM1* are shared with hepatoblasts but together are isolation markers for this population |
| Hepatoblast | *ICAM1*, *KRT8*, *KRT18*, *KRT19* | (Schmelzer *et al*, 2007; Turner *et al*, 2011; So *et al*, 2020; Lemaigre, 2009) | *ICAM1* up-regulation marks the stem-cell-to hepatoblast transition. *KRT8*/*KRT18* are hepatocyte-lineage cytokeratins and *KRT19* a biliary-lineage cytokeratin; biopotential hepatoblasts/progenitors co-express before lineage commitment |
| Hepatocyte | *AFP*, *ALB*, *CEBPA*, *HNF4A*, *MAT1A* | (Lemaigre, 2009; Si-Tayeb *et al*, 2010; Ramani & Lu, 2017; Zong & Stanger, 2012; Camper & Tilghman, 1989; Yeo *et al*, 2024) | *AFP* is characteristic of foetal/early hepatocytes; shared enhancers with *ALB* refocus activity from *AFP* to *ALB* as development proceeds. *CEBPA*/*HNF4A* are core hepatocyte-identity transcription factors that |

| | | | |
|---|---|---|---|
| | | | repress the biliary program. *MAT1A* is adult-hepatocyte-specific (methionine metabolism). |

**Appendix Table S2**: Summary of canonical liver development markers used for annotation of expression clusters.

| Patient | Timepoint 1 (T1) | Timepoint 2 (T2) |
|---|---|---|
| Patient 1 (P1) | 11785 (0.0%) | 73 (0.0%) |
| Patient 2 (P2) | 23921 (0.0%) | 13237 (0.8%) |

**Appendix Table S3**: Post-filtering cell numbers by patient and timepoint. Values shown are post-filtering cell counts, with the percentage of cells removed during QC shown in brackets.

| Cluster ID | Cell Type | InferCNV gains | InferCNV losses | Wesley et al. (2022) | HB Signatures | Development markers |
|---|---|---|---|---|---|---|
| P1_1 | Mostly hepatocytes | 1p, 1q, 20, 16q | 2, 8, 18q | Hepatocyte, (weak) endothelial | 14q32, (weak) C1 | (Strong) *ALB*, *CEBPA*, *AFP*, *EPCAM*, *ICAM1*; (weak) *PROM1* |
| P1_2 | Hepatocytes | 1p, 1q, 20, 16q | 2, 8, 18q | Hepatocyte | 14q32, C1 | (Strong) *MAT1A*, *HNF4A*, *EPCAM*, *KRT8*, *KRT18*; (weak) *ALB*, *CEBPA* |
| P1_3 | Mostly hepatocytes | 1p, 1q, 20, 16q | 2, 8, 18q | Hepatocyte | 14q32, (weak) C1 | (Strong) *ICAM1*, *NCAM1*; (weak) *ALB*, *AFP*, *KRT8*, *KRT18*, *EPCAM* |
| P1_4 | Hepatocytes | 1p, 1q, 20, 16q | 2, 8, 18q | Hepatocyte | C1 | *HNF4A*, *KRT8*, *KRT18*, *KRT19*, |

| | | | | | | |
|---|---|---|---|---|---|---|
| | | | | | | *PROM1*, *CLDN3* |
| P1_5 | Mostly hepatocytes, immune cells | (Weak signal) 20 | (Weak signal) 8, 18q | InferCNV, SingleR annotation, and UMAP segregation consistent with significant stroma proportion in these clusters | | |
| P1_6 | SMCs, TSCs, osteoblasts | 11q | None | | | |
| P1_7 | Mostly SMCs | 11q | None | | | |
| P1_8 | Mostly endothelial | (Weak signal) 1p, 20, 11q | (Weak signal) 2, 8 | | | |
| P1_9 | Macrophages, monocytes, DCs | None | None | | | |
| P2_1 | Hepatocytes | 1q, 20 | 18q | Hepatocyte | 14q32; (weak) C2 | *ALB*, *AFP*, *MAT1A*, *CLDN3* |
| P2_2 | Hepatocytes | 1q, 20 | 18q | Hepatocyte | 14q32; (weak) C2 | *ALB*, *AFP*, *EPCAM*, *ICAM1*, *KRT19* |
| P2_3 | Hepatocytes | 1q, 2, 8, 20 | 18q | Hepatocyte | 14q32; (weak) C2 | (Weak) *ALB*, *AFP*, *MAT1A*, *KRT8*, *KRT18* |
| P2_4 | Mostly hepatocytes | 1q, 2, 20 | 18q | Hepatocyte | C2; (weak) 14q32, Epi-CB, C2 | (Weak) *ALB*, *AFP*, *HNF4A*, *CEBPA*, *KRT8*, *KRT19* |
| P2_5 | Hepatocytes | 1q, 2, 20 | 18q | Hepatocyte | (Weak) 14q32, Epi-CB, C2 | (Weak) *HNF4A*, *CEBPA*, *EPCAM*, *KRT8*, *KRT18*, *KRT19* |
| P2_6 | Mostly hepatocytes | 1q, 20 | 18q | Hepatocyte | (Weak) 14q32, C2 | (Weak) *AFP*, *MAT1A*, *NCAM1*, *ICAM1* |
| P2_7 | Mostly hepatocytes | 1q, 2, 20 | 18q | Hepatocyte | (Weak) 14q32, C2 | (Weak) *HNF4A*, *CEBPA*, *KRT8*, *KRT19*, *NCAM1* |

| | | | | | | |
|---|---|---|---|---|---|---|
| P2_8 | Hepatocytes | 1q, 2, 20 | 18q | (Weak) hepatocyte, cholangiocyte | C1 | *HNF4A*, *CEBPA*, *KRT8*, *KRT18*, *PROM1* |
| P2_9 | Hepatocytes, endothelial | (Weak signal) 1q | None | InferCNV, SingleR annotation, and UMAP segregation consistent with significant stroma proportion in this cluster | | |

**Appendix Table S4**: Cluster annotation table for clustering of clusters identified per patient. Cell type was annotated using SingleR; InferCNV was run to suggest CNV gains and losses; custom annotation was based on the expression of signatures and genes representative of liver development obtained from literature ; and the expression of HB signatures from literature[11,15,25] was also assessed.

| HuH6 MuTrans attractor well depths | | | |
|---|---|---|---|
| *Pre-treatment* | | *Post-treatment* | |
| **Well** | **Depth** | **Well** | **Depth** |
| 0 | 0.243 | 0 | 0.209 |
| 1 | 0.449 | 1 | 0.113 |
| 2 | 0.0867 | 2 | 0.116 |
| 3 | 0.0785 | 3 | 0.108 |
| 4 | 0.148 | 4 | 0.170 |
| | | 5 | 0.285 |

**Appendix Table S5**: HuH6 attractor well depths derived from MuTrans analysis on scRNA-seq data. Larger values for depth correspond to greater stability, i.e. represent a more stable cellular state.

| HepG2 MuTrans attractor well depths | | | |
|---|---|---|---|
| *Pre-treatment* | | *Post-treatment* | |
| **Well** | **Depth** | **Well** | **Depth** |
| 0 | 0.456 | 0 | 0.136 |
| 1 | 0.140 | 1 | 0.176 |
| 2 | 0.133 | 2 | 0.227 |
| 3 | 0.0252 | 3 | 0.136 |
| 4 | 0.116 | 4 | 0.247 |

| | | | |
|---|---|---|---|
| 5 | 0.130 | 5 | 0.0782 |

**Appendix Table S6**: HepG2 attractor well depths derived from MuTrans analysis on scRNA-seq data. Larger values for depth correspond to greater stability, i.e. represent a more stable cellular state.

| Cell line | Non-targeting control (NTC) | *BIRC5* siRNA |
|---|---|---|
| HuH6 | 5531 (2.3%) | 5195 (1.2%) |
| HepG2 | 8044 (1.1%) | 153 (0.0%) |
| HB303 | 8835 (0.1%) | 2753 (0.0%) |

**Appendix Table S7**: Post-filtering cell numbers by cell line and condition (NTC, *BIRC5* siRNA knockdown). Values shown are post-filtering cell counts, with the percentage of cells removed during QC filtering shown in brackets.

Camper SA & Tilghman SM (1989) Postnatal repression of the alpha-fetoprotein gene is enhancer independent. *Genes Dev* 3: 537–546

Chen D, Zhan Y, Zhou E, Chen H, Dai S, Yang R, Ye C, Wang J, Li Y, Lu Y, *et al* (2025) Single-cell profiles delineate immune cell remodeling and enhanced tumor-fibroblast interaction of hepatoblastoma after chemotherapy. *Cell Rep Med* 6: 102401

Lemaigre FP (2009) Mechanisms of Liver Development: Concepts for Understanding Liver Disorders and Design of Novel Therapies. *Gastroenterology* 137: 62–79

Ramani K & Lu SC (2017) Methionine adenosyltransferases in liver health and diseases. *Liver Res* 1: 103–111

Schmelzer E, Zhang L, Bruce A, Wauthier E, Ludlow J, Yao H, Moss N, Melhem A, McClelland R, Turner W, *et al* (2007) Human hepatic stem cells from fetal and postnatal donors. *J Exp Med* 204: 1973–1987

Si-Tayeb K, Lemaigre FP & Duncan SA (2010) Organogenesis and Development of the Liver. *Dev Cell* 18: 175–189

So J, Kim A, Lee S-H & Shin D (2020) Liver progenitor cell-driven liver regeneration. *Exp Mol Med* 52: 1230–1238

Turner R, Lozoya O, Wang Y, Cardinale V, Gaudio E, Alpini G, Mendel G, Wauthier E, Barbier C, Alvaro D, *et al* (2011) Human Hepatic Stem Cell and Maturational Liver Lineage Biology. *Hepatology* 53: 1035–1045

Yeo YH, Lee Y-T, Tseng H-R, Zhu Y, You S, Agopian VG & Yang JD (2024) Alpha-fetoprotein: Past, present, and future. *Hepatol Commun* 8

Zong Y & Stanger BZ (2012) Molecular mechanisms of liver and bile duct development. *WIREs Dev Biol* 1: 643–655
